# Non-autonomy of age-related morphological changes in the *C. elegans* germline stem cell niche

**DOI:** 10.1101/2025.06.13.658151

**Authors:** Nilay Gupta, Mia Sinks, E. Jane Albert Hubbard

## Abstract

Declines in tissue renewal and repair due to alterations in tissue stem cells is a hallmark of aging. Many stem cell pools are maintained morphologically complex niches. Using the *C. elegans* hermaphrodite germline stem cell system, we analyzed age-related changes in the morphology of the niche, the distal tip cell (DTC), and identified a molecular mechanism that promotes a subset of these changes. We found decreases in the number and length of long DTC processes with age. We also found that a long-lived *daf-2* mutant exhibits a *daf-16*-dependent maintenance of long DTC processes. Surprisingly, the tissue requirement for *daf-16(+)* is non-autonomous, and *daf-16(+)* in body wall muscle is both necessary and sufficient. In addition, after a delay, pre-formed DTC processes deteriorate upon premature germline differentiation, but not upon cell cycle inhibition. We propose a reciprocal DTC-germline interaction model and speculate how reduced *daf-2* activity both delays stem cell exhaustion and maintains DTC processes. These studies establish the *C. elegans* DTC as a powerful *in vivo* model for understanding age-related changes in cellular morphology and their consequences in stem cell systems.

**SUMMARY:** The *C. elegans* germline stem cell niche morphology is markedly altered with age and is regulated non-autonomously from the muscle by insulin/IGF-like signaling. Results suggest reciprocal niche-germline regulation.

## INTRODUCTION

Age-related changes in cell morphology can alter cell-cell interactions and signaling (Lopez-Otin et al., 2023). Such changes are particularly important to understand in stem cell systems where niche cells can take on complex morphologies and associate intimately with the stem cells they regulate (Brunet et al., 2023). Age-related changes in the cytoskeleton, extracellular matrix, cell adhesion properties, and other factors influence cell morphology and function (DiLoreto and Murphy, 2015).

Here, we investigate age-related morphological changes in a simple stem cell niche *in vivo*. Like many mammalian stem cell systems (Brunet et al., 2023; Lopez-Otin et al., 2023), the *C. elegans* hermaphrodite germline stem cell pool becomes depleted with age (Garigan et al., 2002; Killian and Hubbard, 2005; Kocsisova et al., 2019; Luo et al., 2010; Qin and Hubbard, 2015). The niche in this system is a single cell, the distal tip cell (DTC), that caps the end of the gonad (Fig. 1) and produces membrane-bound DSL family ligands, APX-1 and LAG-2, that activate Notch pathway signaling in juxtaposed germline stem cells (Austin and Kimble, 1987; Henderson et al., 1994; Kimble and White, 1981; Nadarajan et al., 2009). The adult DTC morphology is complex, with variable numbers and lengths of long cytoplasmic processes that extend over and between germline stem and progenitor cells (Byrd et al., 2014; Hall et al., 1999; Tolkin et al., 2024). DTC processes are not essential for establishment of the stem cell pool that occurs during larval stages, but they contribute to contact-dependent Notch signaling in adults (Lee et al., 2016).

**Figure 1.**
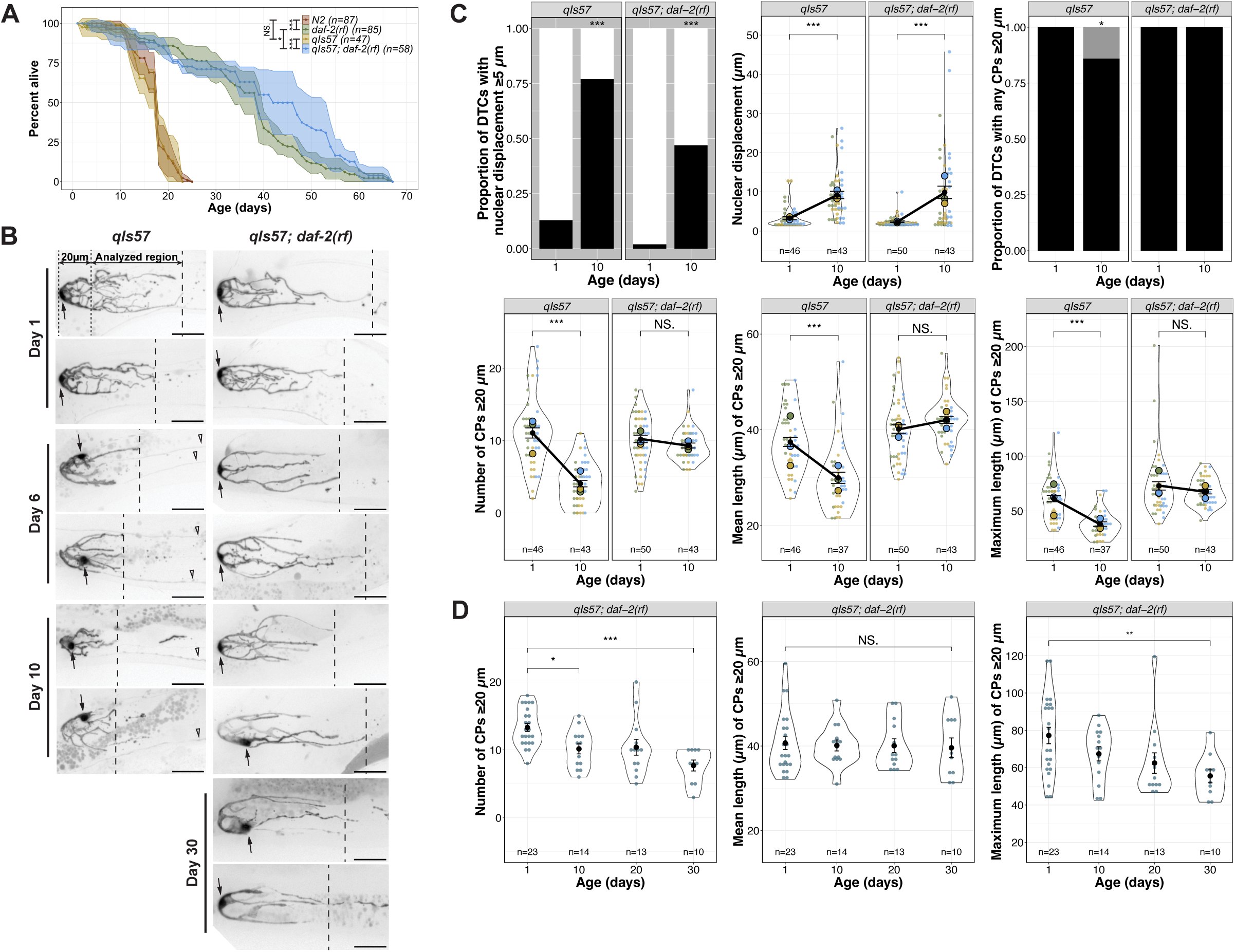
The age-related shift in DTC nuclear position and decline in DTC processes number and length have different levels of dependence on *daf-2*. **(A)** Survival curves for *daf-2(+)* and *daf-2(rf)* worms with and without the *qIs57*[*lag-2p::GFP*] transgene. N2 is the standard wild type, and age is indicated in days post mid-L4 stage. **(B)** Representative micrographs of DTCs in live worms of the indicated genotypes at adult Day 1, 6 and 10, post mid-L4. The 20 µm region is between the dotted lines, and the area analyzed for number and length of continuous processes (CPs; see Gupta et al., 2024; Tolkin et al., 2024) is indicated between the dotted and dashed lines, the latter showing the position of the longest CP. Arrows indicate the DTC nucleus. Open arrowheads indicate non-DTC structures. Scale bars are 20 µm. **(C)** Quantified age-related DTC features (from top left to bottom right): proportion of DTCs with nuclear displacement ≥5 µm (black bar), position of nuclear displacement (µm) from the distal end, proportion of DTCs with any CPs ≥20 µm (black bar), number of CPs ≥20 µm per DTC, mean and maximum length of CPs ≥20 µm per DTC. N values for proportion plots are the same as for the nuclear displacement superplot (top middle). For all superplots, each dot represents a single DTC (n value shown), colors indicate parallel cohorts distinct from those in other main figures, large circles in color are averages for each cohort, the black dot is the pooled average for all cohorts shown, and black line connects pooled average values, indicating the direction and extent of differences between Day 1 and Day 10. Day 10 n values for mean and maximum CP length are lower since only DTCs with CPs >20 µm were included in these measurements. **(D)** Number, mean and maximum length of CPs in *daf-2(rf)* at indicated ages in days post mid-L4. **For all panels**, *qIs57* carries *lag-2p::GFP* and *daf-2(rf)* is *daf-2(e1370).* See methods, Tables S2, S3 for statistics details; comparisons are made between Day 1 and Day 10 within a given genotype; for proportion plots, only p < 0.05 are indicated; otherwise, NS is “not significant”, or p ≥ 0.05; * p < 0.05; ** p < 0.01; *** p < 0.001.

Here we present an aging time-course analysis of changes in DTC morphology and identify molecular mechanisms that delay or hasten them. Our analysis reveals several striking changes to DTC morphology with age. We confirm an age-related proximal shift in nuclear position that occurs at high penetrance (as seen by others under different life history and age conditions (Kocsisova et al., 2019; Urman et al., 2024)), and that the number and length of long continuous DTC processes decline with age. We examined these phenotypes for their dependence on the DAF-2 insulin-like signaling (IIS) pathway, which influences lifespan. IIS had a relatively minor effect on the nuclear position, but the effects of age on DTC process number and length were highly IIS-dependent. In the *daf-2* mutant, long DTC processes persist with age, and this persistence requires DAF-16/FOXO. Surprisingly, the cellular requirement for *daf-16* is neither the DTC itself, nor does it appear to be the intestine where *daf-16* is implicated in regulating longevity. Instead, *daf-16(+)* activity in the body wall muscle is both necessary and sufficient to regulate age-related changes to the length of the DTC processes. We also determined that pre-formed DTC processes deteriorate prematurely by differentiating the underlying germ line, but not by impeding cell cycle. We propose a reciprocal DTC-germline interaction feedback model whereby DTC processes contribute to stem cell maintenance while stem cells stabilize DTC processes, supporting mutual maintenance in young adults (and in *daf-2* mutants) and mutual collapse with age.

## RESULTS

### Adult DTC morphology is altered with age

Age-related changes to hermaphrodite DTC features have been reported previously on a short aging timeline and/or in mated worms (to 5 days of adulthood). These include a proximally displaced nucleus (Kocsisova et al., 2019; Urman et al., 2024) and an increase in the size, though not number, of gaps seen using a membrane reporter (Urman et al., 2024). We sought to further characterize changes to DTC morphology in intact live self-fertile worms over a more extensive timeline from Day 1 to Day 10 of adulthood, and to define morphometric parameters to describe and compare age-related changes under different genetic conditions.

To analyze DTC morphology over time, we selected three time points starting from the mid-L4 as determined by vulval morphology (Mok et al., 2015): Day 1 (24 hours post mid-L4), when DTC elaboration has peaked in the wild type; Day 6 post mid-L4, when the reproductive phase of wild-type self-fertile hermaphrodites is ending; and Day 10 post mid-L4, when worms are on the verge of the steep population decline consistent with their ∼2-3 week lifespan. To minimize variability due to subtle differences in rearing conditions, for each cohort analyzed, worms were selected as mid-L4 larvae on the same day and harvested for live imaging on Days 1, 6, and 10.

We examined several different transgenes expressing fluorescent proteins in the DTC, settling on *qIs57* which encodes GFP under the control of the *lag-2* promoter (Siegfried et al., 2004). Compared to the other transgenes we examined, it revealed a suitably detailed set of morphological features, did not appreciably dim with age, and allowed tracking of the position of the nucleus (Fig. S1). To ensure that the transgene was not itself modulating lifespan, we performed lifespan analyses in parallel with the canonical wild-type strain N2 and found no significant difference (Fig. 1A). However, a greater percentage of worms were censored in strains bearing *qIs57* (see Materials and methods). Adult DTCs are morphologically variable; two DTCs from each timepoint are shown in Figure 1B and additional examples are in Figure S2.

Using our image analysis pipeline (Gupta et al., 2024), we found several striking changes to DTC morphology with age. Day 1 to Day 10 provides the most informative comparisons (Fig. 1B,C; see Fig. S3 for additional timepoints). First, recapitulating and extending the findings of others (Kocsisova et al., 2019; Urman et al., 2024), we observed that the DTC nucleus, which normally sits at the distal end of the DTC, shifts proximally with age. The percentage of DTCs with the nucleus ≥5 µm from the distal end went from 13% at Day 1 to 77% at Day 10, and these nuclei averaged 3 µm and 9 µm from the distal end at Day 1 and Day 10, respectively. Second, the number and length of processes extending proximally from the DTC (Byrd et al., 2014; Hall et al., 1999; Tolkin et al., 2024) provided additional parameters that change with age. LAG-2 fusion proteins can be visualized along DTC processes (Henderson et al., 1992; Crittenden et al., 2006; Gordon et al., 2019) and germ cell proximity to processes is associated with higher probability of signal-responding germ cells (Lee et al., 2016). Therefore, changes to DTC processes could conceivably influence the underlying germ cell fate. Though additional parameters may be relevant, we focused on long DTC processes that appeared continuous with the cell body in maximum projection micrographs (“continuous processes” (CPs); Gupta et al., 2024; Tolkin et al., 2024). Our methods do not distinguish between previously-defined subclasses of DTC processes that intercalate with or extend along the germ cells (Byrd et al., 2014), but they reflect only gross morphological features including the proportion of DTCs with processes that extend beyond a 20 µm threshold from the distal end, and the numbers and lengths (mean and maximum) of such processes. At Day 1 post mid-L4, all DTCs had at least one process extending beyond 20 µm. By Day 10, 14% of DTCs had no CPs beyond this threshold. Among those DTCs with one or more CP over the threshold, we observed fewer and shorter CPs over time: the mean number of CPs decreased from 11 to 4 from Day 1 to Day 10, as did their mean and maximum lengths (mean 37 to 30 µm, and maximum 60 to 38 µm) (Fig. 1C).

### Insulin/insulin-like growth factor-1 signaling (IIS) promotes age-related changes in DTC process number and length

The IIS pathway is highly conserved and modulates longevity in *C. elegans*, *Drosophila*, and mice (Kenyon, 2010b; Murphy and Hu, 2013; Russell and Kahn, 2007). In addition to longevity, IIS regulates various physiological processes in *C. elegans*, including development, metabolism, and stress resistance (Murphy and Hu, 2013). DAF-2 is the sole insulin/IGF-1 receptor in *C. elegans* (Kimura et al., 1997), and reducing *daf-2* activity (e.g. by the reduction-of-function *(rf) daf-2(e1370)* allele) extends lifespan (Kenyon et al., 1993). To determine whether a reduction in *daf-2* activity would affect age-related changes in DTC morphology, we examined DTCs in *daf-2(rf)* worms. First, we tested whether the *qIs57* marker affected the *daf-2(rf)* longevity phenotype and found that, if anything, the percentage of worms alive between 40-60 days was elevated, but the maximum lifespan was similar (Fig. 1A). We aged cohorts of wild-type and *daf-2(rf)* worms in parallel and measured the parameters as described above.

First, we found that *daf-2(rf)* had a minimal and variable effect on age-related nuclear position displacement (Figs 1C, S3A). Relative to the wild type, *daf-2(rf)* displayed a reduced proportion of DTCs that exhibit nuclear displacement of ≥5 µm at Day 10. However, a large percentage (47%) of *daf-2(rf)* worms nevertheless displayed the nuclear displacement phenotype (versus 77% in the wild type), and the average degree of displacement in microns from the distal end was similar to the wild type in this cohort. The latter finding is consistent with those of Kocsisova et al. (Kocsisova et al., 2025, their Fig. S1). In summary, nuclear displacement occurs at a high penetrance in both wild type and *daf-2(rf)* with an equal distance of displacement, suggesting that IIS plays a minor role in regulating this aspect of age-related changes to DTC morphology.

In contrast to the minor effects on nuclear position, reducing *daf-2* activity virtually halted age-related changes in CP number and length from Day 1 to Day 10: all *daf-2(rf)* worms possessed at least one CP ≥20 µm at Day 10, and there was no decrease in the average number of CPs ≥20 µm, nor their mean or maximum lengths, as was seen in the wild type (Figs 1C, S4). We therefore focused on the DTC phenotypes related to number and length of CPs.

### The persistence of DTC process number and length in daf-2(rf) does not correlate temporally with extended lifespan or reproductive span

In addition to lifespan extension in *daf-2(rf)*, the reproductive span of self-fertile hermaphrodites averages several days longer (Dillin et al., 2002), and in mated hermaphrodites, the *e1370* allele, in particular, delays reproductive aging (Kocsisova et al., 2025). We wondered whether the rate of age-related changes in the number and length of DTC processes might correlate with reproductive span and/or population lifespan. If correlated with reproductive span, we would expect DTC processes to be shorter after a week in *daf-2(rf)*. If the rate of decline were associated with average lifespan, we would expect to observe shorter processes by 30 days. We therefore collected additional measurements in *daf-2(rf)* up to Day 30 (Fig. 1D; additional timepoints in Fig. S3B). Referencing our lifespan measurements, Day 30 in *daf-2(rf)* reflects the percent alive of wild type at Day 15. We found that, remarkably, while the number of CPs declined modestly in this cohort by Day 30 (Fig. 1D), it still remained above that of the wild type at Day 10 (Fig. 1C). In addition, there was no statistically significant decline in the mean length of CPs ≥20 µm up to 30 days. Although we observed a significant decline in the maximum length, the average is still longer than the wild type at Day 10.

We conclude that reducing *daf-2* markedly delays age-related changes in the number and length of long DTC processes, and that these changes in DTC morphology do not directly correlate with temporal extension of reproductive span or of population lifespan.

### Insulin/insulin-like growth factor-1 signaling (IIS) acts via DAF-16/FOXO to promote age-related changes in DTC process length

Many phenotypes of *daf-2(rf)* mutants, including age-related phenotypes such as lifespan, depend on the DAF-16/FOXO transcription factor (Kenyon, 2010a; Murphy and Hu, 2013).To determine whether the *daf-2*-dependent age-related changes in DTC process number and length are also dependent on *daf-16*, we examined DTC morphology, as visualized with the *qIs57* marker, in double mutant strains bearing a *daf-16* null mutation *daf-16(mu86)* and *daf-2(e1370)* [hereafter referred to as *daf-16(0); daf-2(rf)*]. Although the vast majority of DTCs had one or more CP that exceeded 20 µm, the number of processes and their mean and maximum length decreased similar to the wild type (Fig. 2).

**Figure 2.**
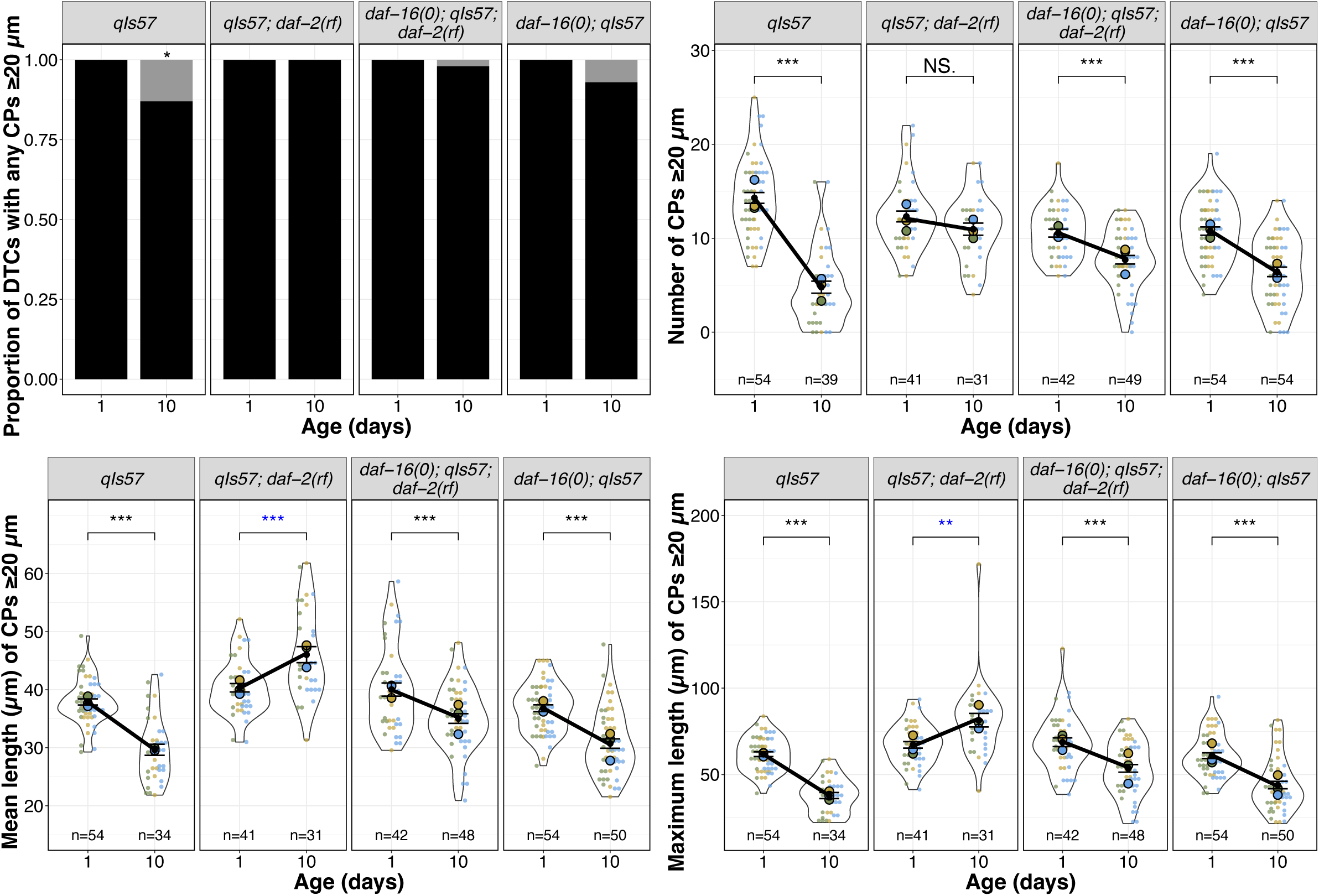
Loss of *daf-16* largely reverses the effects of *daf-2* on CP number and length. From top to bottom: proportion of DTCs with any CPs ≥20 µm (black bars), number of CPs ≥20 µm per DTC, mean and maximum length of CPs ≥20 µm per DTC in the indicated genotypes for worms imaged on Day 1 and Day 10 post mid-L4. N values for proportion plots correspond to the “Number of CPs” superplot to the right; n values for mean and maximum length plots reflect only those DTCs with CPs ≥ 20 µm. For superplots, dots and circles are as described in Figure 1; colors indicate cohorts and replicates are distinct from those presented in other main figures. See methods, Tables S2, S3 for statistics details; NS is “not significant”, * p < 0.05, ** p < 0.01, *** p < 0.001, and blue asterisks indicate a significant increase from Day 1 to Day 10.

To assess the loss of *daf-16* alone, we measured the same parameters in parallel cohorts with the wild type, *daf-2(rf)*, and *daf-16(0); daf-2(rf)* (Figs 2, S5). In these trials, a small percentage of CPs of Day 10 DTCs in *daf-16(0)* worms did not exceed 20 µm, though not as many as the wild type. CP number and length also decreased, as in the wild type. Taken together, we conclude that the maintenance of DTC process number and length over time in *daf-2(rf)* is highly dependent on *daf-16* activity.

### Non-autonomous daf-16 activity influences age-related decline of DTC process length

Prior studies indicated that *daf-16* activity in specific tissues can influence distinct phenotypes while also contributing to systemic effects. For example, intestinal *daf-16* is associated with enhanced stress resistance and increased longevity (Libina et al., 2003; Zhang et al., 2022), and *daf-16* in the germ line and muscle promotes expansion of the progenitor pool during germline development (Michaelson et al., 2010). Additionally, *daf-16* activity in *fos-1a* expressing cells in the proximal somatic gonad (PSG) prevents the age-dependent decline in number of germline progenitors (Qin and Hubbard, 2015). Therefore, we sought to determine in which tissues *daf-16* most influences the maintenance of DTC processes with age that we observed in *daf-2(rf)*. For this analysis, we focused on the DTC length parameters (mean and maximum).

Our first approach took advantage of available extrachromosomal arrays (Libina et al., 2003; Qin and Hubbard, 2015). Each includes a tissue-restricted promoter driving a GFP::DAF-16a translational fusion. We performed the aging time course in the *daf-16(0); daf-2(rf)* background and compared DTC processes length parameters from control and array-bearing (phenotypically Rol) worms, and non-array bearing (phenotypically non-Rol) progeny from the same array-bearing mothers. If *daf-16(+)* activity were important in a particular tissue location, we expected that its expression would preserve DTC process length relative to *daf-16(0); daf-2(rf)*.

To assess the approach, we first examined CPs in worms bearing an array that expresses *daf-16a* from the *daf-16* promoter. We observed that these parameters (the proportion of worms with DTC CPs over 20 µm, and the CP length – both mean and maximum) were markedly stable relative to *daf-16(0); daf-2(rf)* (Fig. 3) and to non-array bearing siblings (Fig. S6). The phenotypic similarity of *daf-16(0); daf-2(rf)* to non-array bearing progeny from array-bearing mothers also suggested that the array did not confer a maternal effect.

**Figure 3.**
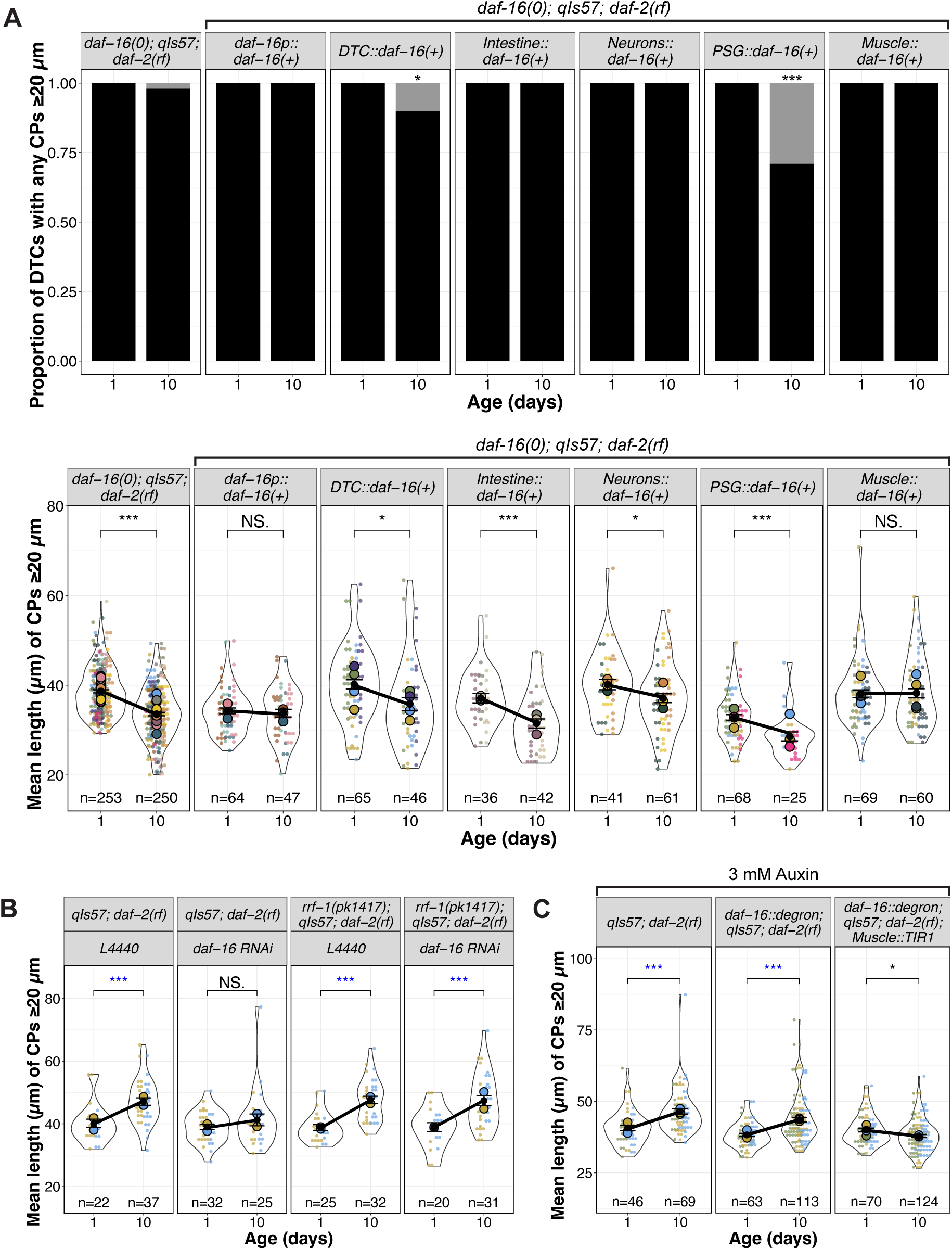
*daf-16(+)* in body wall muscle is sufficient and necessary for *daf-2(rf)* suppression of age-related changes in DTC process length. **(A)** Proportion of DTCs with CPs ≥20 µm, and mean length of CPs ≥20 µm per DTC for indicated genotypes (*daf-16(0); qIs57; daf-2(rf)* alone and with extrachromosomal arrays expressing *daf-16a(+)*) at Day 1 and Day 10 post mid-L4. N values for mean length plots reflect only those DTCs with CPs ≥ 20 µm. See Fig. S6 for additional timepoints and maximum length.**(B)** Superplots showing mean length of CPs ≥20 µm in genotypes indicated. *L4440* indicates the empty vector control and *daf-16 RNAi* is the vector carrying *daf-16* cDNA sequences. See Fig. S8 for additional timepoints and maximum length. **(C)** Mean length of CPs ≥20 µm in worms raised on 3mM auxin in genotypes indicated, where *daf-16::degron* is *hq389[daf-16::GFP::degron]* and *Muscle::TIR1* is *emcSi71[myo-3p::TIR-1::mRuby]*. See Fig. S10 for images, additional time points, and maximum length. **For all panels,** *qIs57* carries *lag-2p::GFP*, *daf-16(0)* is *daf-16(mu86),* and *daf-2(rf)* is *daf-2(e1370).* For all superplots, colors indicate cohorts within panels; replicates shown here are distinct from those in other main figures. See methods, Tables S2, S3 for statistics details; NS is “not significant”, * p < 0.05, ** p < 0.01, *** p < 0.001, and blue asterisks indicate a significant increase.

We next tested for DTC-autonomy, expressing *daf-16a* from the *lag-2* promoter (Michaelson et al., 2010). Surprisingly, ∼15% of ∼140 DTCs exhibited an unusual morphology at Days 6 and 10, half displaying uncharacteristically long DTC processes, while others displayed one thick CP containing a displaced nucleus (Fig. S7). None of these features were observed under any other condition. Among the remaining ∼85% of DTCs that exhibited normal morphology, the CP length parameters were similar to *daf-16(0); daf-2(rf),* even though ∼10% of the DTCs in array-bearing worms at Day 10 had no CPs ≥ 20 µm (Figs 3, S6). Thus, apart from the low-penetrance alterations in gross morphology, *lag-2*-driven *daf-16a(+)* had no appreciable effect on the persistence of DTC process length with age. We conclude that non-autonomous *daf-16a(+)* activity is sufficient to prevent the age-related decline in DTC length when *daf-2* is reduced.

### Intestinal activity of daf-16a does not appear to promote age-related DTC length in daf-2(rf)

Both heterologous expression approaches (Libina et al., 2003) and protein degradation approaches (Zhang et al., 2022) indicate that the intestine is the major contributor of *daf-16* activity that confers extended lifespan in *daf-2(rf)*. We speculated that changes in DTC morphology might be a function of overall organismal aging. If so, we would expect that intestinal *daf-16(+)* in the *daf-16(0); daf-2(rf)* mutant background would stabilize DTC processes, similar to *daf-16(+)* driven by its own promoter.

Again, using the same transgene expressing *daf-16(+)* that alters lifespan (Libina et al., 2003), we tested the sufficiency of *daf-16a* expressed from an intestine-specific promoter. We found that although all worms bore DTCs with CPs ≥ 20 µm, CP lengths still declined significantly with age, comparable to that observed in *daf-16(0); daf-2(rf)* controls (Figs 3, S6). In conclusion, although intestinal *daf-16(+)* contributes the majority of the longevity phenotype in *daf-2(rf)* (Libina et al., 2003; Zhang et al., 2022), it does not appear to regulate the length of DTC processes with age.

### Neuronal daf-16a(+) activity mildly influences the age-related changes in DTC length

Although the intestine is the major contributor of *daf-16* on lifespan, there is neuronal contribution as well (Uno et al., 2021; Zhang et al., 2022). We tested an array expressing *daf-16a(+)* from a pan-neuronal promoter and observed that while all DTCs in the worms bearing the neuronally-expressed *daf-16a(+)* array contained CPs that crossed the 20 µm threshold, the CPs showed a modest but significant length decline with age. We conclude that neuronal activity of *daf-16(+)* makes a minor contribution to maintaining the length of DTC processes with age in *daf-2(rf)*.

### Proximal somatic gonad (PSG) daf-16a(+) activity does not prevent the age-related loss of DTC process length

Previous results indicate that *daf-16a* expressed in *fos-1a-*positive cells of the proximal somatic gonad (PSG) in a *daf-16(0); daf-2(rf)* double mutant delays age-dependent loss of germline progenitors (Qin and Hubbard, 2015), and *daf-16*-dependent DOS-3, a non-canonical ligand for the GLP-1/Notch receptor, is a relevant PSG signal (Zhang et al., 2024). We hypothesized the PSG-expressed *daf-16* might also contribute to DTC morphology. Although prior experiments used *naEx239[*pGC629*(fos1p::gfp::daf-16a) +* pRF4] in *daf-16(m26); daf-2(e1370)* (Qin and Hubbard, 2015), for consistency between experiments here, we moved the array into the *daf-16(mu86); daf-2(rf)* background, and measured CP number and length.

In short, *daf-16a(+)* activity in the PSG did not prevent the age-dependent loss of DTC process length. However, these worms were markedly less healthy than other strains bearing *daf-16(+)* arrays: only ∼50% of worms survived to Day 4, a defect not observed with other *daf-16(+)* arrays. Moreover, nearly a third (29%) of worms bearing the array had no DTC process ≥20 µm. Of those that did, the mean and maximum lengths were lower than the control on Day 1 and nevertheless continued to decrease with age (Figs 3A, S6).

Collectively, these findings demonstrate that the expression of *daf-16a(+)* via *fos-1a* promoter is insufficient to prevent the age-dependent DTC process decline in a *daf-16(0); daf-2(rf)* background. This result raises the possibility that while *daf-16a(+)* in the PSG promotes maintenance of the progenitor pool with age (Qin and Hubbard, 2015; Zhang et al., 2024), it does not delay the decline of DTC process length with age.

### Germline daf-16 exerts a minor effect on DTC process length

The germ line is the major focus of activity for *daf-16* in modulating expansion of the progenitor zone (PZ) in larval stages (Michaelson et al., 2010), and the development of adult DTC morphology is partially dependent on the germ line (Tolkin et al., 2024). To determine whether germline *daf-16* is required for long continuous DTC processes to persist in *daf-2(rf)*, we used RNAi feeding and the *rrf-1* mutant in which RNAi is less effective in the soma but is retained in the germ line (Kumsta and Hansen, 2012; Sijen et al., 2001). We note that in these experiments (as in Fig. 2) *daf-2(rf)* displays an increase in mean (Fig. 3B) and maximum (Fig. S8) CP length from Day 1 to Day 10, an effect that is not observed with *rrf-1* alone (Fig. S8). We speculate that this may be related to a delay in the PZ reaching its maximum size in *daf-2(rf)* worms (see Discussion). As expected from the foregoing mutant analysis, reducing *daf-16* by RNAi eliminated the mean and maximum CP length increase observed in *daf-2(rf)* from Day 1 to Day 10. In the absence of *rrf-1*, however, the mean CP length increased in both the L4440 control and with *daf-16(RNAi)*, suggesting that the contribution of *daf-16* from the soma is more relevant for mean CP length. For maximum length (Fig. S8), a minor effect of *daf-16(RNAi)* is seen in the absence of *rrf-1*. We conclude that germline *daf-16* plays a minor to negligible role in regulating DTC processes in *daf-2(rf)* with age.

### Body wall muscle-specific daf-16a(+) is sufficient and necessary for the DTC process length reduction with age in daf-2(rf)

We measured DTC parameters in *daf-16(0); daf-2(rf)* worms bearing an array expressing *daf-16a(+)* in body wall muscle (*myo-3p::GFP::daf-16a*; Libina et al., 2003). All the worms carrying the array had CPs ≥20 µm (Fig. 3A). Furthermore, CP length was maintained in *daf-16(0); daf-2(rf)* relative to controls without the array and was similar to worms carrying the *daf-16p::daf-16(+)* array (Figs 3A, S6). We also did not observe a significant effect of muscle *daf-16(+)* on the extension of reproductive span that is seen in *daf-2(rf)* alone nor an increase in self-fertile brood size (Fig. S9A). Together, these findings indicate that array-borne expression of *daf-16a* in body wall muscle is sufficient to maintain the length of DTC processes with age in *daf-2(rf)*.

To assess the necessity of body wall muscle DAF-16, we used an auxin-mediated degron approach (Zhang et al., 2015). We generated a strain bearing *daf-2(rf),* a *TIR1::mRub*y fusion expressed under the *myo-3* promoter (Sabatella et al., 2021) together with degron-tagged DAF-16::GFP (Zhang et al., 2022), and the DTC marker. Following the red and green fluorescent protein tags, we confirmed the tissue-specific expression of TIR1 and auxin-dependent loss of DAF-16 (Fig. S10A). In all of the vehicle controls (*daf-2(rf)* and the DAF-16::degron alone), the DTCs in *daf-2(rf)* strains not only maintained but increased in length over the 10 day interval, while mean length of DTC processes in the strain bearing both *myo-3p::TIR1* and the degron-tagged DAF-16 decreased (Figs 3C, S6). Although DAF-16::GFP was undetectable with auxin exposure, the relatively modest effect could be due to residual undetectable DAF-16.

These results suggest that *daf-16* in body wall muscle is a major contributor to the persistence of long DTC processes in *daf-2(rf)* with age.

### Body wall muscle DAF-16a attenuates the rate of germline progenitor zone decline with age

Prior studies implicated muscle-expressed *daf-16a* activity in a minor but significant role downstream of *daf-2* in promoting larval expansion of the germline progenitor pool (Michaelson et al., 2010), whereas it did not significantly influence the endpoint of the progenitor cell number in aged worms (Qin and Hubbard, 2015).

Our DTC morphology analysis prompted us to re-examine the possibility of a role for *myo-3p::daf-16a(+)* in maintaining the PZ pool over an aging time-course in *daf-2(rf)*. We found that the PZ pool of *daf-16(0); daf-2(rf)* worms expressing *myo-3p::daf-16a(+)* started with fewer cells than *daf-16(0); daf-2(rf)* on Day 1, consistent with the results of Michaelson et al. (2010). Relative to controls, *daf-16(0); daf-2(rf)* with *myo-3p::daf-16a(+)* also does not affect the terminal PZ cell count, consistent with the results of Qin and Hubbard (2015). However, the presence of the *myo-3p::daf-16a(+)* array does attenuate the rate of decline of the PZ, similar to *daf-2(rf)* (Figs 4A, S9B). This result suggests that the slower PZ loss over time in *daf-2(rf)* is partially dependent on muscle-produced *daf-16(+)*. Other tissues likely contribute, since an array bearing *daf-16(+)* driven from its own promoter gives a stronger effect. By contrast, muscle-produced *daf-16(+)* prevents loss of DTC processes length over time, similar to what is observed with *daf-16(+)* driven by its own promoter.

**Figure 4.**
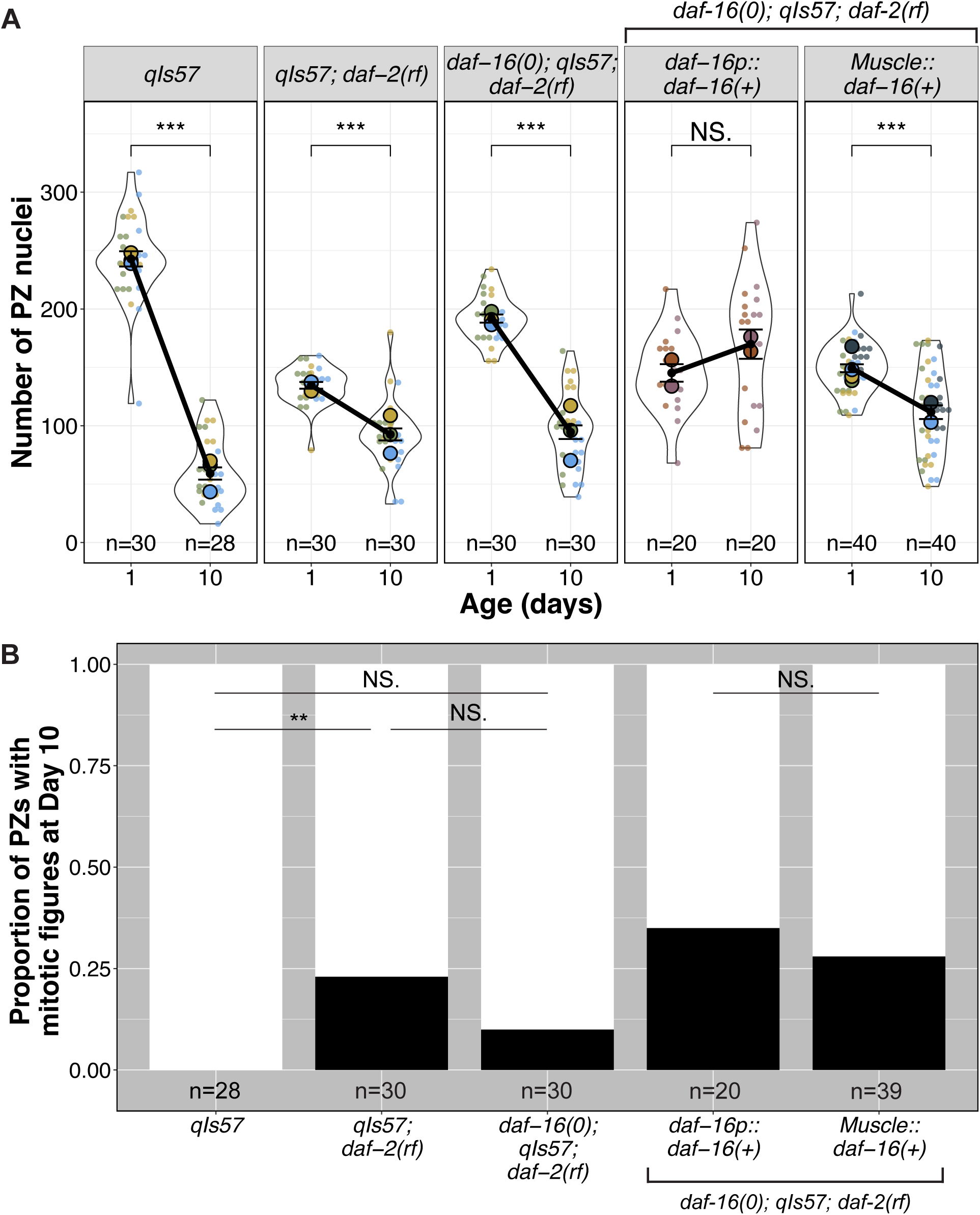
Muscle-expressed *daf-16* influences both the rate of progenitor zone (PZ) pool loss and continued germ cell divisions with age. **(A)** The number of PZ nuclei in (n) gonads in worms of the indicated genotypes at Day 1 and Day 10 post mid-L4. **(B)** The proportion of PZs in Day 10 adults displaying one or more mitotic figures. See Fig. S9C for mitotic index. See Table S3 for all pairwise comparisons. **For all panels,** *qIs57* carries *lag-2p::GFP*, *daf-16(0)* is *daf-16(mu86),* and *daf-2(rf)* is *daf-2(e1370).* For all superplots, colors indicate cohorts within panels. See methods, Tables S2, S3 for statistics details; NS is “not significant”, *** p < 0.001.

We further asked whether altering *daf-2* and *daf-16,* and in particular *daf-16a(+)* from muscle, would influence the proportion of gonads displaying active cell divisions (mitotic figures) on Day 10, a time-point at which we observed no mitotic figures in the wild type. We found significantly more *daf-2(rf)* worms display mitotic figures at Day 10, while *daf-16(0); daf-2(rf)* do not, though the *daf-16(0); daf-2(rf)* proportion is intermediate, suggesting that only part of this phenotype may be *daf-16-*dependent. Of *daf-16(0); daf-2(rf)* worms expressing *myo-3p::daf-16a(+)*, 28% display mitotic figures, similar to worms expressing *daf-16p::daf-16a(+)*. These results suggest that muscle-expressed *daf-16a(+)* could play a role in maintaining Day 10 germ cell cycling in self-fertile *daf-2(rf)* hermaphrodites. Overall, muscle-expressed *daf-16(+)* both delays age-related changes to the PZ pool and to the DTC, with more subtle effects on the PZ than the DTC, though other tissues also contribute to each independently (see Discussion).

### Germline undifferentiated fate status, but not cell cycle progression, is correlated with maintenance of DTC processes

A striking feature of the aged *daf-2(rf)* germ line is the relative stability of the PZ pool compared to the wild type (Figs 4A, S9B). In mated worms, *daf-2(rf)* also displays a more proximal border of SYGL-1-positive stem cells and a more constant position of meiotic entry relative to the wild type (Kocsisova et al., 2025). We further observed that the proportion of gonads with active divisions in Day 10 is higher in *daf-2(rf)* and the mitotic index did not decline significantly from Day 1 to Day 10, and that all these phenotypes are partially dependent on *daf-16* (Figs 4A-B, S9B-C). These results prompted us to consider the relationship between DTC processes, stem cell fate, and mitotic progression.

Out-growth of the DTC processes in early adulthood is dependent on an underlying substrate of the undifferentiated germ cells (Byrd et al., 2014; Tolkin et al., 2024). We therefore wondered whether adult DTC processes, once formed, would be stable in the absence of the underlying PZ in early adulthood. We analyzed the number and length of long DTC processes in worms bearing a temperature sensitive allele of *glp-1/* Notch, *glp-1(e2141),* in which the entire PZ differentiates at the restrictive temperature (Austin and Kimble, 1987). We examined the nuclear morphology of distal germ cells and measured DTC parameters from parallel cohorts of worms that underwent the same temperature-shift regime (Fig. 5A). After 24 hours at the restrictive temperature, although virtually all distal germ cells had entered meiotic prophase, the CP number and length parameters were unaffected (Fig. S11A-C). By 48 hours, however, the CP length was reduced relative to the wild type (Fig. 5A-C). We conclude that premature differentiation of the PZ pool causes a premature decline in the complexity of the DTC morphology, but with a time delay between differentiation and the decline of DTC processes.

**Figure 5.**
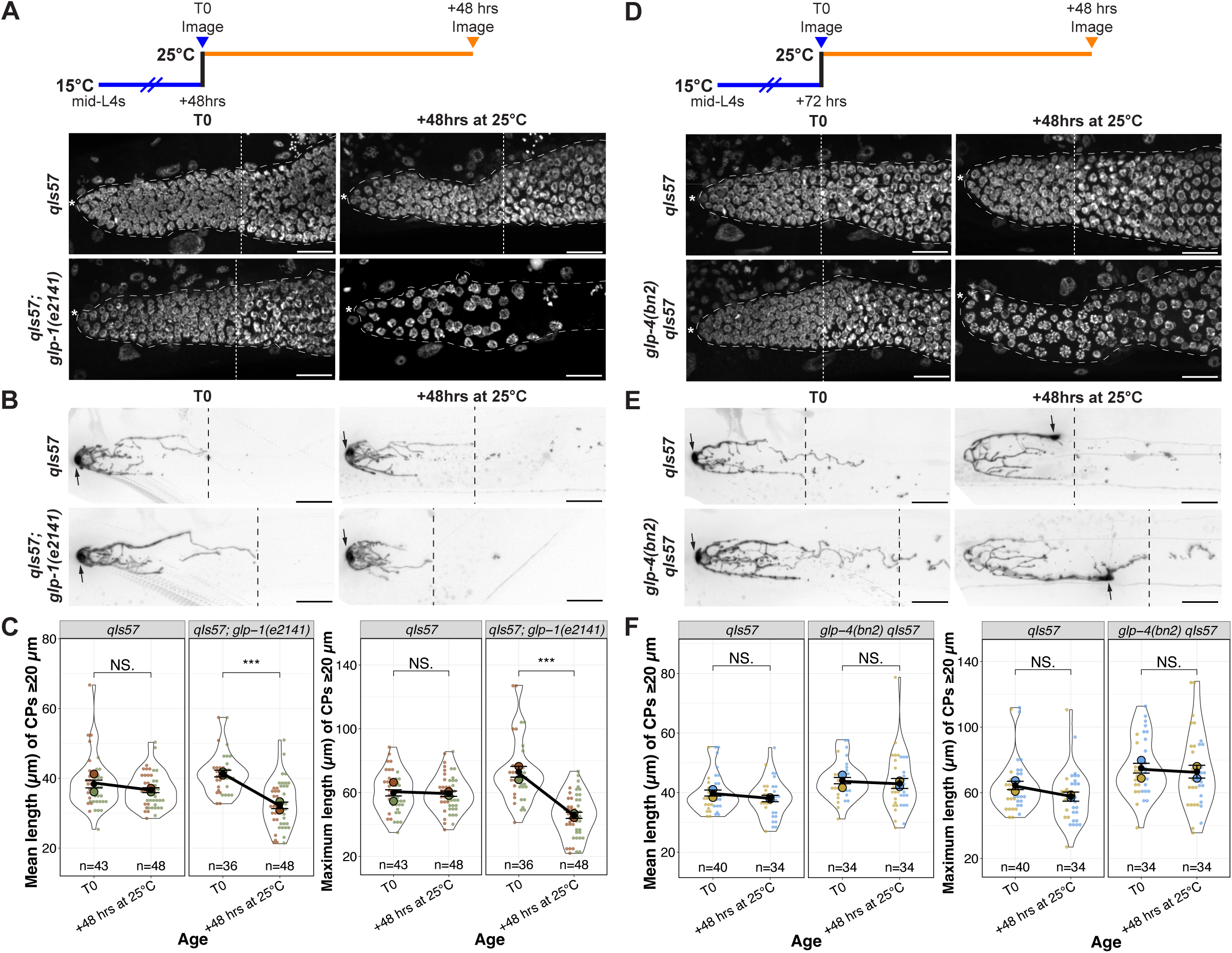
DTC processes decline prematurely with germline stem cell differentiation, but not with restricted cell cycle progression. **(A, D)** Experimental design and images of DAPI-stained control and *glp-1(e2141ts)* (A) or *glp-4(bn2ts)* (D) worms at 48 hours after shift to the restrictive temperature. White dotted lines indicate the proximal border of the PZ. Asterisks indicate the distal end. **(B, E)** Examples of DTC morphology in worms scored in parallel to those shown in panel (A). Black dashed lines indicate the longest CP. Arrows indicate the DTC nucleus. **(A,B,D,E)** Scale bars are 20 µm. **(C, F)** Quantification of mean and maximum lengths of CPs ≥20 µm in control and *glp-1(e2141ts)* (C) or *glp-4(bn2ts)* (F) worms at indicated time points. Colors indicate cohorts within panels. See methods, Tables S2, S3 for statistics details; NS is “not significant”, * p < 0.05, *** p < 0.001.

**Figure 6.**
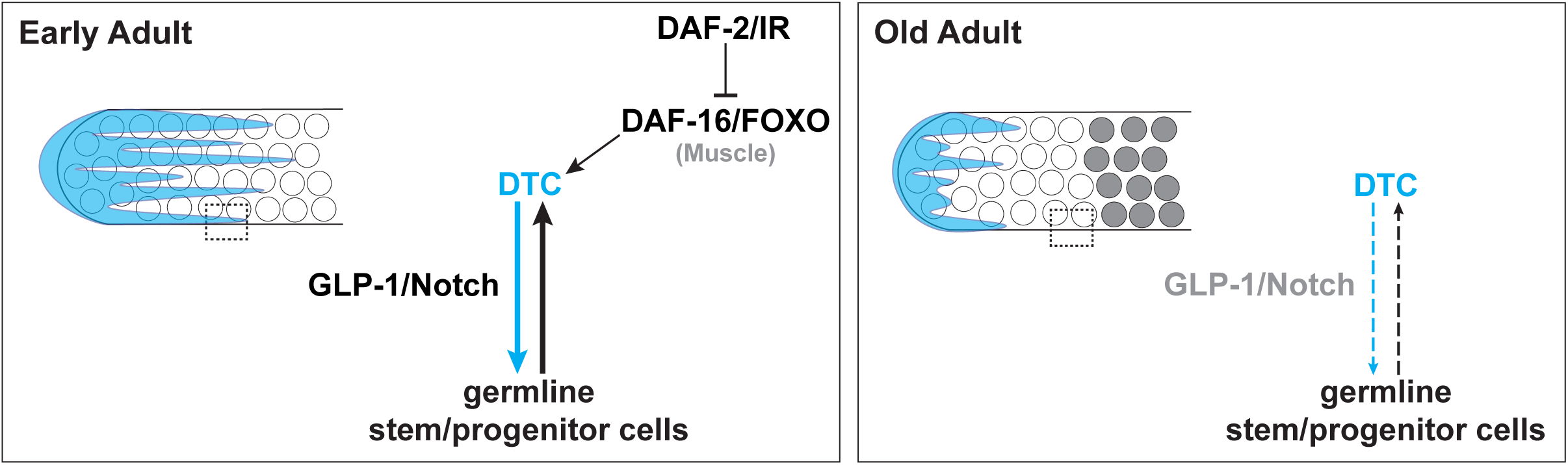
Cartoon representation of a two-component feedback model for niche-stem cell interaction at early and late (“old”) adult stages. Dotted box indicates area of direct DTC-germline interaction that is lost with age. White circles represent undifferentiated germ cells, and grey circles represent differentiated germ cells. See text for details.

Because differentiated germ cells no longer divide, and because *daf-2(rf)* display more mitotic figures than the wild type at a late time point (Day 10; Figs 4B, S9C), and this phenotype is partially dependent on muscle-expressed *daf-16(+)*, we wondered whether the DTC process maintenance requires active cycling in the underlying germline stem/progenitor cells. To test this possibility, we used a temperature sensitive allele of *glp-4, glp-4(bn2),* that causes germ cell cycle arrest in pro-metaphase; arrested *glp-4* germ cells are resistant to differentiation, even in the absence of *glp-1* (Beanan and Strome, 1992). Remarkably, after 48 hours at the restrictive temperature, although *glp-4* mutant gonads display altered nuclear morphology, the DTC processes did not collapse (Figs 5D-F, S12A-C). We further observed that germ cells were capable of recovery after shifting back to the permissive temperature for another 48 hours, and the DTC processes remained intact (Fig S12A-C).

Therefore, while loss of stem cells to differentiation eventually leads to changes in DTC morphology that resemble what is seen in older adults, interfering with cell cycle without causing differentiation (as in *glp-4* at the restrictive temperature) does not.

## DISCUSSION

Here we define age-related changes in DTC niche morphology and assess their dependence on IIS, a molecular mechanism that affects longevity. We found that an age-related shift in DTC nucleus position is only modestly affected by IIS in self-fertile hermaphrodites, as was previously shown in mated worms (Kocsisova et al., 2019), while the decline in the number and length of long DTC processes is highly dependent on IIS. We identify body wall muscle as a major source of a *daf-16(+)*-dependent activity that can support both the persistence of long DTC processes in a *daf-2(rf)* mutant and the underlying germline progenitor pool. Finally, we determine a dependency relationship between the germ cell proliferative state and maintenance of pre-formed DTC processes.

### A model for reciprocal feedback interactions between the aging germline and niche

In a seeming paradox, *daf-2(+)* expands the larval germline progenitor zone (PZ) but drives loss of the PZ with age, and both effects are dependent on *daf-16*. In larvae, the germline cell cycle is slower in worms with reduced *daf-2* activity and this is dependent on *daf-16(+)* in the germ line and body wall muscle (Michaelson et al., 2010; Roy et al., 2016). In aged adult worms, a larger PZ pool is maintained with reduced *daf-2* activity (Qin and Hubbard, 2015; Fig. 4A). In this case, one *daf-16*-dependent signal, DOS-3, from the proximal somatic gonad likely supports GLP-1/Notch activity in the germ line (Zhang et al., 2024). We found that, in addition, both the PZ pool and long DTC processes are maintained in old *daf-2(rf)* adults in a *daf-16(+)* muscle-autonomous manner. The exact *daf-16*-dependent mechanism – and whether it acts on the directly on the DTC, the germline, or both – awaits further study. We speculate that such a factor could be a *daf-16*-dependent secreted factor – e.g., a mitokine or myokine signaling mechanism, or some other mechanism secondary to the IIS role in muscle function (Herndon et al., 2002; Wang et al., 2019). There is precedent for body wall muscle-produced secreted proteins acting on the germ line, including HIM-4/hemicentin (Vogel and Hedgecock, 2001) and SWM-1 (Chavez et al., 2018).

Regardless of the exact mechanism, these observations suggest a model for interplay between the control of cell fate and cell cycle rate that may contribute to the dynamics of the germline stem cell system over time with age. Several additional relevant features of the system are noted below.

First, stem cell fate and cell cycle rate are independently controlled. The PZ consists of a distal-most SYGL-1(+) germline stem cells (GSCs) that are actively responding to GLP-1/Notch (Kershner et al., 2013), as well as their more proximal SYGL-1(-) progeny that complete a cell division before differentiating (Fox and Schedl, 2015). Reducing *glp-1* activity moves the border of the PZ distally and reduces the number of GSCs but, importantly, does not alter their rate of cell cycle progression (Fox and Schedl, 2015; Michaelson et al., 2010; Roy et al., 2016). By contrast, reducing *daf-2* slows the cell cycle (e.g., lowers M-phase and S-phase indices, a rough proxy) in larvae but does not influence the stem versus non-stem cell fate decision governed by GLP-1/Notch (Michaelson et al., 2010; Roy et al., 2016). Worms with reduced *daf-2* activity therefore reach adulthood with fewer cells PZ cells. This pool declines with age, but at a rate far slower than the wild type (Kocsisova et al., 2022; Qin and Hubbard, 2015; Fig. 4A). Second, as in many mammalian stem cell systems (Brunet et al., 2023), *C. elegans* GSCs become depleted over time (Kocsisova et al., 2019; Qin and Hubbard, 2015), and, importantly, the PZ pool is effectively “used up” with age. This loss is accelerated in the presence of replete sperm that encourage germline flux (Kocsisova et al., 2019; Qin and Hubbard, 2015). In addition, with age, the mitotic index slows, even in the presence of replete sperm (Kocsisova et al., 2019). Finally, in early adulthood, the outgrowth of DTC processes partially depends on adhesion to underlying proliferative germ cells (Tolkin et al., 2024).

Taking these observations together, we propose a two-component feedback system that supports homeostasis of the progenitor pool in early adulthood, and the loss of which contributes to the collapse of both niche morphology and the stem cell pool with age. In this model, the early adult PZ steady state is maintained by a combination of DTC-to-GSC signaling via GLP-1/Notch to designate the stem cell fate in responding germ cells, as well as DAF-2-dependent inputs that promote continued stability of DTC processes. Over time, however, as cells leave the PZ through differentiation (meiotic entry) and the pool of undifferentiated germ cells becomes depleted, overlying DTC processes are not maintained due to both loss of DAF-16-dependent muscle (and other) effects, as well as PZ depletion. Decreased ligand availability due to shortened DTC processes would then further deplete the stem cell pool and further destabilize the DTC processes.

In the *daf-2(rf)* scenario, we propose that in addition to the effect on *daf-16*, the persistence of slow-cycling yet undifferentiated germline stem/progenitor cells may contribute to the stability of DTC processes, perhaps by adhesion mechanisms similar to those that act in the early adult (Tolkin et al., 2024). This stability could then support DTC-to-GSC interactions along the DTC processes that help maintain stem cell fate (Fig. 5D). Again, GLP-1/Notch signaling afforded by stable DTC processes may, in turn, support germ cells in an undifferentiated, proliferation-competent (and continued slow-cycling) state. Reducing *daf-2* would thereby delay the age-related declines in both components of the reciprocal interaction that are mutually supportive, the progenitor pool and niche morphology.

Temporally, the decline of the stem cell pool and PZ precedes that of the DTC processes (Figs 1, 4). Similarly, when germ cells are forced to differentiate, a time delay occurs before the DTC declines (Fig. 5). We observed that loss of *glp-1*, which causes germ cells in the PZ to differentiate, leads to premature collapse of already-formed DTC processes while loss of *glp-4*, which impedes the cell cycle but does not cause differentiation does not cause the collapse of DTC processes. These results implicate the undifferentiated state of germ cells in the germline-to-DTC branch of the feedback to help maintain DTC processes.

While this model provides a framework for reciprocal signaling between DTC processes and the germ line, additional influences act on the system. For example, the regulation of GLP-1/Notch signaling with age (Kocsisova et al., 2019) is complex: in addition to DOS-3-mediated signaling from the proximal somatic gonad (Zhang et al., 2024), the shift in the position of the DTC nucleus also alters the patterning of signaling (Urman et al., 2024), and there may be germline-autonomous aspects of reduced GLP-1/Notch signaling since, at least to Day 4, *sygl-1* RNA levels do not change (Urman et al., 2024). In mated worms, reduced Notch signaling is implicated in depletion of the stem cell pool and reproductive aging, both of which are also delayed in mated *daf-2(rf)* worms (Kocsisova et al., 2025). It will be of interest to determine, ideally in individual gonads, to what extent reduced ligand signaling due to changes in DTC morphology with age contribute to the age-related decline in the germline stem cell pool and subsequent reproductive aging.

Our model also predicts that the effects of DAF-16 from the proximal somatic gonad that act on the germ line should affect the DTC too, via feedback, since they preserve the PZ. However, we did not observe the predicted stabilization of DTC processes when *daf-16(+)* was expressed in the proximal somatic gonad in *daf-16(0); daf-2(rf)*. This result, together with the more marked effect of muscle-expressed *daf-16* on the DTC versus the PZ suggest that the germ cell interaction is not the sole arbiter of DTC stability with age. Finally, many aspects of the germ cell cycle are unusual, including a highly abbreviated G1, constitutively high levels of cyclin E/CDK2 throughout the cell cycle, and transcriptional regulation of CDK-2 through inhibition of DPL-1 (Fox et al., 2011; Furuta et al., 2018). How such cell cycle features are modified with age and how such modifications influence germ cell propensity to differentiate are unresolved.

### General implications for aging stem cell systems and regenerative medicine

Our results demonstrate that niche morphology is a key aspect of aging stem cell systems. Non-autonomous control of niche morphology, by tissues outside the stem cell system and/or the adjacent stem cells, may be important in other stem cell systems as well. Our work further suggests that deeper knowledge of *in vivo* interactions, especially those influenced by cellular morphology and cell cycle as well as cell-cell signaling, are required to understand stem cell behavior and tissue regeneration, especially in aging systems.

## MATERIALS AND METHODS

### Nematode strains and maintenance

The *C. elegans* strains utilized in this study were raised on standard nematode growth media (NGM) agar and *E. coli* OP50 at 20°C, unless otherwise specified. All strains were derived from Bristol N2 (Brenner, 1974). All strains, including their complete genotypes and sources, are provided in Table S1. The strains generated in this study were obtained through genetic crosses and homozygosity was confirmed by progeny testing – visually for strains bearing fluorescent markers or by DNA sequence analysis based on information in WormBase (Davis et al., 2022). Strains generated for this study are GC1593, GC1607, GC1616, GC1676, GC1682, GC1683, GC1691, GC1692, GC1696, GC1748, GC1778, GC1817, GC1819, and GC1912, and are available upon request.

### Aging and lifespan assays

Multiple plates of approximately 20 L4 hermaphrodites per plate were prepared, and worms were transferred daily to fresh plates to prevent overcrowding and mixing with their progeny. Worms were collected for imaging or PZ counting at intervals indicated. Controls were run in parallel within each cohort; cohorts share a color in each figure that reports DTC morphometrics or PZ nuclei counts, or mitotic index. Lifespan assays (Fig. 1A) started with 100 worms per genotype. For N2 and CB1370, ∼15% were censored due to bagging, desiccating on the petri dish wall, or were otherwise missing. For the two strains that contain *qIs57* (JK2869 and GC1607) the rate of censoring for the same reasons was higher, at 40-50%.

### Imaging

With the exception of images of GC1412 in Fig. S1, DTC images were acquired using a Apo 60X oil immersion lens on a Nikon W1 spinning disk confocal microscope and subsequently processed, quantified, and analyzed as described in Gupta et al., 2024 (Gupta et al., 2024). The DTC was visualized using the *qIs57* GFP marker, with a 488 nm excitation laser set at ≤10% power. The exposure time was set to 100 ms or 1 frame, the EM Gain Multiplier was set to 50, the conversion gain was set to 1, and the readout mode was set to ‘EM Gain 10 MHz at 16-bit’. The images were acquired as Z-stacks at 0.5 µm step size and saved in .nd2 format. The excitation laser used for DAPI was 405 nm at 20-30% power, and for RFP was 561 nm at 20-30% power range.

### DAPI staining, progenitor zone (PZ) counts, and mitotic figures and index

DAPI staining (using Vectashield from Vector Laboratories™ H-1200-10) was performed as described in Pepper et al., 2003, except that fixation and staining were done in a 9-well glass depression plates (Corning™ 7220-85) to minimize the loss of older worms. Images were acquired using a 60X oil immersion lens on a Nikon W1 spinning disk confocal microscope. The progenitor zone (PZ) is defined as the region before the occurrence of more than one crescent-shaped nucleus within the same transverse section of the gonad (Crittenden et al., 2006; Killian and Hubbard, 2005). After visualizing the entire stack, the PZ region of interest was identified, and the nuclei were counted manually using the multi-point tool in ImageJ™. Mitotic figures were defined as obvious metaphase, anaphase, and early telophase figures as visualized in images after whole-mount DAPI staining. Mitotic index is the number of mitotic figures over total number of PZ nuclei.

### Auxin-mediated degradation

For auxin-inducible protein degradation (AID) experiments, 3 mM of Indole-3-acetic acid (IAA, auxin) (Thermo Scientific™ A10556.14) or an equivalent volume of carrier EtOH for controls was added to NGM agar prior to pouring plates. OP50 bacteria were concentrated approximately 2x prior to seeding the auxin-containing plates to compensate for auxin inhibition of bacterial growth (Zhang et al., 2022). Worms were cultured on both the auxin-containing plates and the control plates for multiple generations prior to initiating the aging protocol.

### Analysis of glp-1(e2141) and glp-4(bn2)

Worms (*glp-1* and *glp-4* mutant with parallel controls) were collected at the mid-L4 stage and cultured at 15°C for 48 and 72 hours, respectively, to allow for DTC elaboration comparable to that seen in Day 1 adults raised at 20°C. For both genotypes, a group of worms were separated from the cohort and imaged (T0). For *glp-1* experiments, the rest were shifted to 25°C and imaged 24 and 48 hours later. For *glp-4* experiments, the rest were shifted to 25°C for 48 hours and then shifted back to 15°C for another 48 hours. Cohorts of worms that were grown under the same conditions were split and analyzed live for DTC features or for cell counts after DAPI staining.

### RNAi bacterial feeding

RNAi feeding was carried out as previously described (Timmons et al., 2001) with minor modifications. Single colonies of *E. coli* strain HT115 carrying either an empty vector (L4440) or a vector containing *daf-16* cDNA sequences were obtained from frozen stocks on solid LB plates supplemented with 100 µg/ml ampicillin and 10 µg/ml tetracycline, grown overnight at 37°C and used to inoculate liquid LB cultures supplemented with 100 µg/ml ampicillin. Cultures were grown at 37°C with shaking for 6-8 hours and used to seed NGM plates supplemented with 100 µg/ml ampicillin and 0.5% β-lactose. Seeded RNAi plates were kept at room temperature for 24 hours prior to adding worms. Complete penetrance of the *bli-3* Bli phenotype was used as a positive control to confirm the efficacy of RNAi reagents in each experimental replicate. Fresh plates were seeded twice a week over the course of the experiment. L4 hermaphrodites were cultured on either L4440-or *daf-16* RNAi-seeded plates for 1 generation prior to starting the aging assays.

### Statistics

Table S2 (.xlsx file) provides properties analyzed, means, SE/SD, and n values for all data presented in main and supplementary figures (tab for each main and corresponding supplemental figure). Table S3 (.xlsx file) lists the genotypes compared, statistical tests, and the exact p values for each main and supplementary data figure on a separate tab. For lifespan assays, the Mantel-Cox log-rank test was used. Fisher’s exact test was used for pair-wise comparisons of proportion plots (DTCs with nuclear displacement ≥5 µm, DTCs with any CPs ≥20 µm, and PZs with mitotic figures at Day 10). For determining significance between nuclear displacement, number, mean length, and maximum length of CPs ≥20 µm, the number of PZ nuclei, or mitotic index: two-tailed t-test for pairwise comparisons (main figures), and two-tailed t-test with Bonferroni correction for multiple comparisons (supplement). When reporting the mean and maximum CP length, we did not include DTCs that with no CPs >20 µm. All sample numbers (n) are also reported within figures.

## ACKNOWLEDGEMENTS

We thank the CGC, which is funded by NIH Office of Research Infrastructure Programs (P40 OD010440), NYU Langone’s Microscopy Laboratory (RRID: SCR_017934), especially Michael Cammer; this shared resource is partially supported by the Cancer Center Support Grant P30CA016087, Sophia Heimbrock for technical assistance, and Anke Kloock, Theadora Tolkin, Julie Manikas and Caroline Goutte for advice and support.

## FUNDING

NIH R01AG065672

NYSTEM DOH 01-C32560GG-3450000

## Notes

### Competing Interest Statement

The authors have declared no competing interest.

### Summary of Updates

Edits were made to the text and figures, especially to figure 5 and new figure 6.

## REFERENCES

1. Austin, J. and Kimble, J. (1987). glp-1 is required in the germ line for regulation of the decision between mitosis and meiosis in C. elegans. Cell 51, 589–599.

2. Beanan, M. J., & Strome, S. (1992). Characterization of a germ-line proliferation mutation in C. elegans. *Development (Cambridge*, England), 116, 755–766.

3. Brenner, S. (1974). The genetics of Caenorhabditis elegans. Genetics 77, 71–94.

4. Brunet, A., Goodell, M. A. and Rando, T. A. (2023). Ageing and rejuvenation of tissue stem cells and their niches. Nat Rev Mol Cell Biol 24, 45–62.

5. Byrd, D. T., Knobel, K., Affeldt, K., Crittenden, S. L. and Kimble, J. (2014). A DTC niche plexus surrounds the germline stem cell pool in Caenorhabditis elegans. PLoS One 9, e88372.

6. Chavez, D. R., Snow, A. K., Smith, J. R., & Stanfield, G. M. (2018). Soma-germ line interactions and a role for muscle in the regulation of *C. elegans* sperm motility. *Development (Cambridge*, England), 145.

7. Crittenden, S. L., Leonhard, K. A., Byrd, D. T. and Kimble, J. (2006). Cellular analyses of the mitotic region in the Caenorhabditis elegans adult germ line. Molecular biology of the cell 17, 3051–3061.

8. Davis, P., Zarowiecki, M., Arnaboldi, V., Becerra, A., Cain, S., Chan, J., Chen, W. J., Cho, J., da Veiga Beltrame, E., Diamantakis, S., et al. (2022). WormBase in 2022-data, processes, and tools for analyzing Caenorhabditis elegans. Genetics 220.

9. Dillin, A., Crawford, D. K. and Kenyon, C. (2002). Timing requirements for insulin/IGF-1 signaling in C. elegans. Science 298, 830–834.

10. DiLoreto, R. and Murphy, C. T. (2015). The cell biology of aging. Mol Biol Cell 26, 4524–4531.

11. Fox, P. M. and Schedl, T. (2015). Analysis of Germline Stem Cell Differentiation Following Loss of GLP-1 Notch Activity in Caenorhabditis elegans. Genetics 201, 167–184.

12. Fox, P. M., Vought, V. E., Hanazawa, M., Lee, M.-H., Maine, E. M. and Schedl, T. (2011). Cyclin E and CDK-2 regulate proliferative cell fate and cell cycle progression in the C. elegans germline. *Development (Cambridge*, England) 138, 2223–2234.

13. Furuta, T., Joo, H. J., Trimmer, K. A., Chen, S. Y. and Arur, S. (2018). GSK-3 promotes S-phase entry and progression in C. elegans germline stem cells to maintain tissue output. Development 145.

14. Garigan, D., Hsu, A. L., Fraser, A. G., Kamath, R. S., Ahringer, J. and Kenyon, C. (2002). Genetic analysis of tissue aging in Caenorhabditis elegans: a role for heat-shock factor and bacterial proliferation. Genetics 161, 1101–1112.

15. Gordon, K. L., Payne, S. G., Linden-High, L. M., Pani, A. M., Goldstein, B., Hubbard, E. J. A., & Sherwood, D. R. (2019). Ectopic Germ Cells Can Induce Niche-like Enwrapment by Neighboring Body Wall Muscle. Current biology : CB, 29, 823–833.e5.

16. Gupta, N., Cammer, M., Tolkin, T. and Hubbard, E. J. A. (2024). A DTC morphometrics package for quantification of complex and variable cellular morphology using ImageJ. MicroPubl Biol 2024.

17. Hall, D. H., Winfrey, V. P., Blaeuer, G., Hoffman, L. H., Furuta, T., Rose, K. L., Hobert, O. and Greenstein, D. (1999). Ultrastructural features of the adult hermaphrodite gonad of Caenorhabditis elegans: relations between the germ line and soma. Dev Biol 212, 101–123.

18. Henderson, S. T., Gao, D., Lambie, E. J. and Kimble, J. (1994). lag-2 may encode a signaling ligand for the GLP-1 and LIN-12 receptors of C. elegans. *Development (Cambridge*, England) 120, 2913–2924.

19. Herndon, L. A., Schmeissner, P. J., Dudaronek, J. M., Brown, P. A., Listner, K. M., Sakano, Y., Paupard, M. C., Hall, D. H. and Driscoll, M. (2002). Stochastic and genetic factors influence tissue-specific decline in ageing C. elegans. Nature 419, 808–814.

20. Kenyon, C. (2010a). The first long-lived mutants: discovery of the insulin/IGF-1 pathway for ageing. Philosophical Transactions of the Royal Society B: Biological Sciences 366, 9–16.

21. Kenyon, C., Chang, J., Gensch, E., Rudner, A. and Tabtiang, R. (1993). A C. elegans mutant that lives twice as long as wild type. Nature 366, 461–464.

22. Kenyon, C. J. (2010b). The genetics of ageing. Nature 464, 504–512.

23. Kershner, A., Crittenden, S. L., Friend, K., Sorensen, E. B., Porter, D. F. and Kimble, J. (2013). Germline Stem Cells and Their Regulation in the Nematode Caenorhabditis elegans. pp. 29–46. Dordrecht: Springer Netherlands.

24. Killian, D. J. and Hubbard, E. J. A. (2005). Caenorhabditis elegans germline patterning requires coordinated development of the somatic gonadal sheath and the germ line. Developmental Biology 279, 322–335.

25. Kimble, J. E. and White, J. G. (1981). On the control of germ cell development in Caenorhabditis elegans. Dev Biol 81, 208–219.

26. Kimura, K. D., Tissenbaum, H. A., Liu, Y. and Ruvkun, G. (1997). daf-2, an insulin receptor-like gene that regulates longevity and diapause in Caenorhabditis elegans. Science 277, 942–946.

27. Kocsisova, Z., Bagatelas, E. D., Santiago-Borges, J., Lei, H. C., Egan, B. M., Mosley, M. C., Anderson, A. M., Schneider, D. L., Schedl, T., & Kornfeld, K. (2025). Notch signaling in germ line stem cells controls reproductive aging in *Caenorhabditis elegans*. PNAS nexus, 4.

28. Kocsisova, Z., Kornfeld, K. and Schedl, T. (2019). Rapid population-wide declines in stem cell number and activity during reproductive aging in C. elegans. Development 146.

29. Kumsta, C. and Hansen, M. (2012). C. elegans rrf-1 mutations maintain RNAi efficiency in the soma in addition to the germline. PLoS One 7, e35428.

30. Lee, C., Sorensen, E. B., Lynch, T. R. and Kimble, J. (2016). C. elegans GLP-1/Notch activates transcription in a probability gradient across the germline stem cell pool. eLife 5.

31. Libina, N., Berman, J. R. and Kenyon, C. (2003). Tissue-specific activities of C. elegans DAF-16 in the regulation of lifespan. Cell 115, 489–502.

32. Lopez-Otin, C., Blasco, M. A., Partridge, L., Serrano, M. and Kroemer, G. (2023). Hallmarks of aging: An expanding universe. Cell 186, 243–278.

33. Luo, S., Kleemann, G. A., Ashraf, J. M., Shaw, W. M. and Murphy, C. T. (2010). TGF-β and insulin signaling regulate reproductive aging via oocyte and germline quality maintenance. Cell 143, 299–312.

34. Michaelson, D., Korta, D. Z., Capua, Y. and Hubbard, E. J. A. (2010). Insulin signaling promotes germline proliferation in C. elegans. *Development (Cambridge*, England) 137, 671–680.

35. Mok, D. Z., Sternberg, P. W. and Inoue, T. (2015). Morphologically defined sub-stages of C. elegans vulval development in the fourth larval stage. BMC Dev Biol 15, 26.

36. Murphy, C. T. and Hu, P. J. (2013). Insulin/insulin-like growth factor signaling in C. elegans. WormBook, 1–43.

37. Nadarajan, S., Govindan, J. A., McGovern, M., Hubbard, E. J. A. and Greenstein, D. (2009). MSP and GLP-1/Notch signaling coordinately regulate actomyosin-dependent cytoplasmic streaming and oocyte growth in C. elegans. *Development (Cambridge*, England) 136, 2223–2234.

38. Qin, Z. and Hubbard, E. J. A. (2015). Non-autonomous DAF-16/FOXO activity antagonizes age-related loss of C. elegans germline stem/progenitor cells. Nature Communications 6, 7107.

39. Roy, D., Michaelson, D., Hochman, T., Santella, A., Bao, Z., Goldberg, J. D. and Hubbard, E. J. A. (2016). Cell cycle features of C. elegans germline stem/progenitor cells vary temporally and spatially. Dev Biol 409, 261–271.

40. Russell, S. J. and Kahn, C. R. (2007). Endocrine regulation of ageing. Nat Rev Mol Cell Biol 8, 681–691.

41. Sabatella, M., Thijssen, K. L., Davo-Martinez, C., Vermeulen, W. and Lans, H. (2021). Tissue-Specific DNA Repair Activity of ERCC-1/XPF-1. Cell Rep 34, 108608.

42. Siegfried, K. R., Kidd, A. R., 3rd, Chesney, M. A. and Kimble, J. (2004). The sys-1 and sys-3 genes cooperate with Wnt signaling to establish the proximal-distal axis of the Caenorhabditis elegans gonad. Genetics 166, 171–186.

43. Sijen, T., Fleenor, J., Simmer, F., Thijssen, K. L., Parrish, S., Timmons, L., Plasterk, R. H. and Fire, A. (2001). On the role of RNA amplification in dsRNA-triggered gene silencing. Cell 107, 465–476.

44. Timmons, L., Court, D. L. and Fire, A. (2001). Ingestion of bacterially expressed dsRNAs can produce specific and potent genetic interference in Caenorhabditis elegans. Gene 263, 103–112.

45. Tolkin, T., Burnett, J. and Hubbard, E. J. A. (2024). A role for organ level dynamics in morphogenesis of the C. elegans hermaphrodite distal tip cell. Development 151.

46. Uno, M., Tani, Y., Nono, M., Okabe, E., Kishimoto, S., Takahashi, C., Abe, R., Kurihara, T. and Nishida, E. (2021). Neuronal DAF-16-to-intestinal DAF-16 communication underlies organismal lifespan extension in C. elegans. iScience 24, 102706.

47. Urman, M. A., John, N. S., Jung, T. and Lee, C. (2024). Aging disrupts spatiotemporal regulation of germline stem cells and niche integrity. Biol Open 13.

48. Vogel, B. E., & Hedgecock, E. M. (2001). Hemicentin, a conserved extracellular member of the immunoglobulin superfamily, organizes epithelial and other cell attachments into oriented line-shaped junctions. *Development (Cambridge*, England), 128, 883–894.

49. Wang, H., Webster, P., Chen, L. and Fisher, A. L. (2019). Cell-autonomous and non-autonomous roles of daf-16 in muscle function and mitochondrial capacity in aging C. elegans. Aging (Albany NY) 11, 2295–2311.

50. Zhang, L., Ward, J. D., Cheng, Z. and Dernburg, A. F. (2015). The auxin-inducible degradation (AID) system enables versatile conditional protein depletion in C. elegans. Development 142, 4374–4384.

51. Zhang, Y. P., Zhang, W. H., Zhang, P., Li, Q., Sun, Y., Wang, J. W., Zhang, S. O., Cai, T., Zhan, C. and Dong, M. Q. (2022). Intestine-specific removal of DAF-2 nearly doubles lifespan in Caenorhabditis elegans with little fitness cost. Nat Commun 13, 6339.

52. Zhang, Z., Yang, H., Fang, L., Zhao, G., Xiang, J., Zheng, J. C. and Qin, Z. (2024). DOS-3 mediates cell-non-autonomous DAF-16/FOXO activity in antagonizing age-related loss of C. elegans germline stem/progenitor cells. Nat Commun 15, 4904.

