## Supplemental Figures S1-S12 and Table S1 for "Non-autonomy of age-related morphological changes in the *C. elegans* germline stem cell niche"

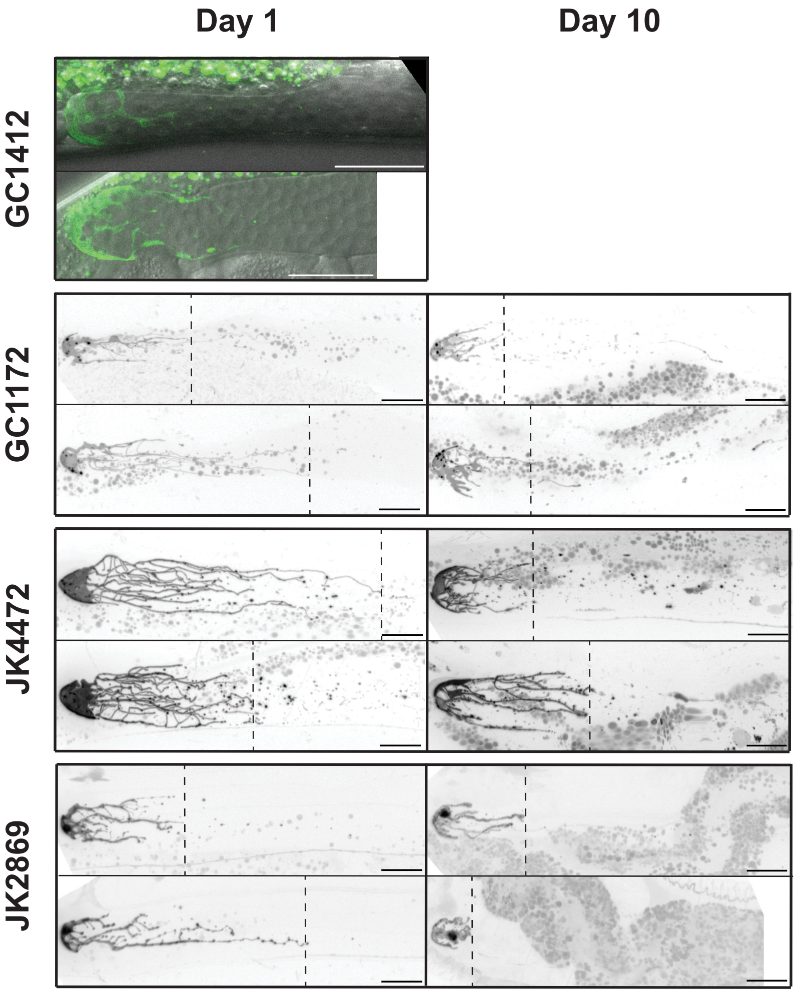


| Strain | Relevant Genotype | Relevant Features |
| --- | --- | --- |
| GC1412 | ***naSi8***<br>Single copy insertion of *lag-2p::GFP-PH* | Dim expression; nucleus not visible |
| GC1172 | ***naIs37***<br>Low-copy insertion of *lag-2p::PH-mCherry* | *mCherry* aggregates; dim expression; nucleus not visible |
| JK4472 | ***qIs154***<br>Insertion of *lag-2p::Myr-tdTomato* | Strong localization to the plasma membrane; variable intensity; high detail; nucleus not visible |
| JK2869 | ***qIs57***<br>Insertion of *lag-2p::GFP* | Strong fluorescence signal throughout the cell; high and consistent intensity; moderate detail; clearly visible nucleus |

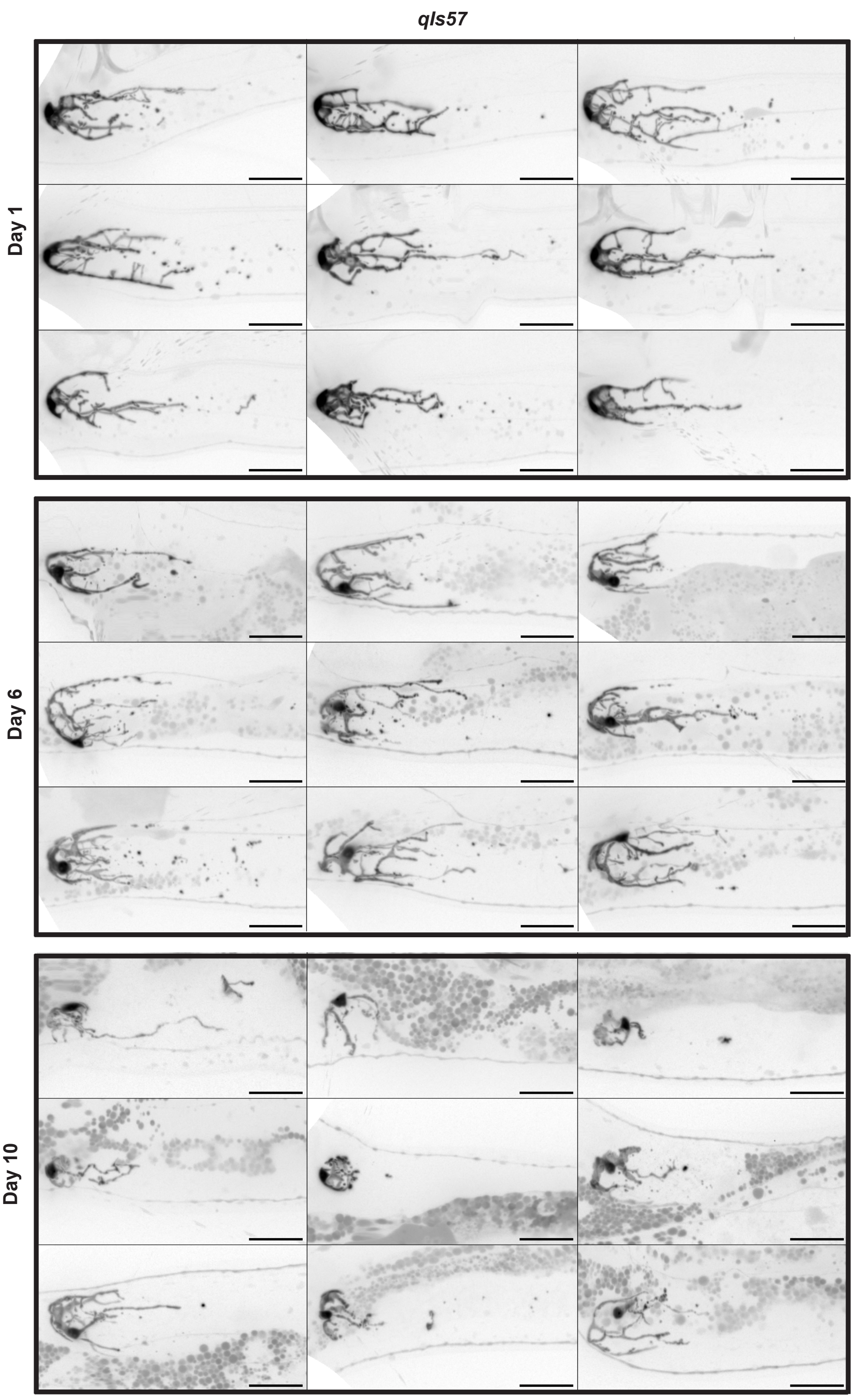
qIs57
Day 1
Day 6
Day 10

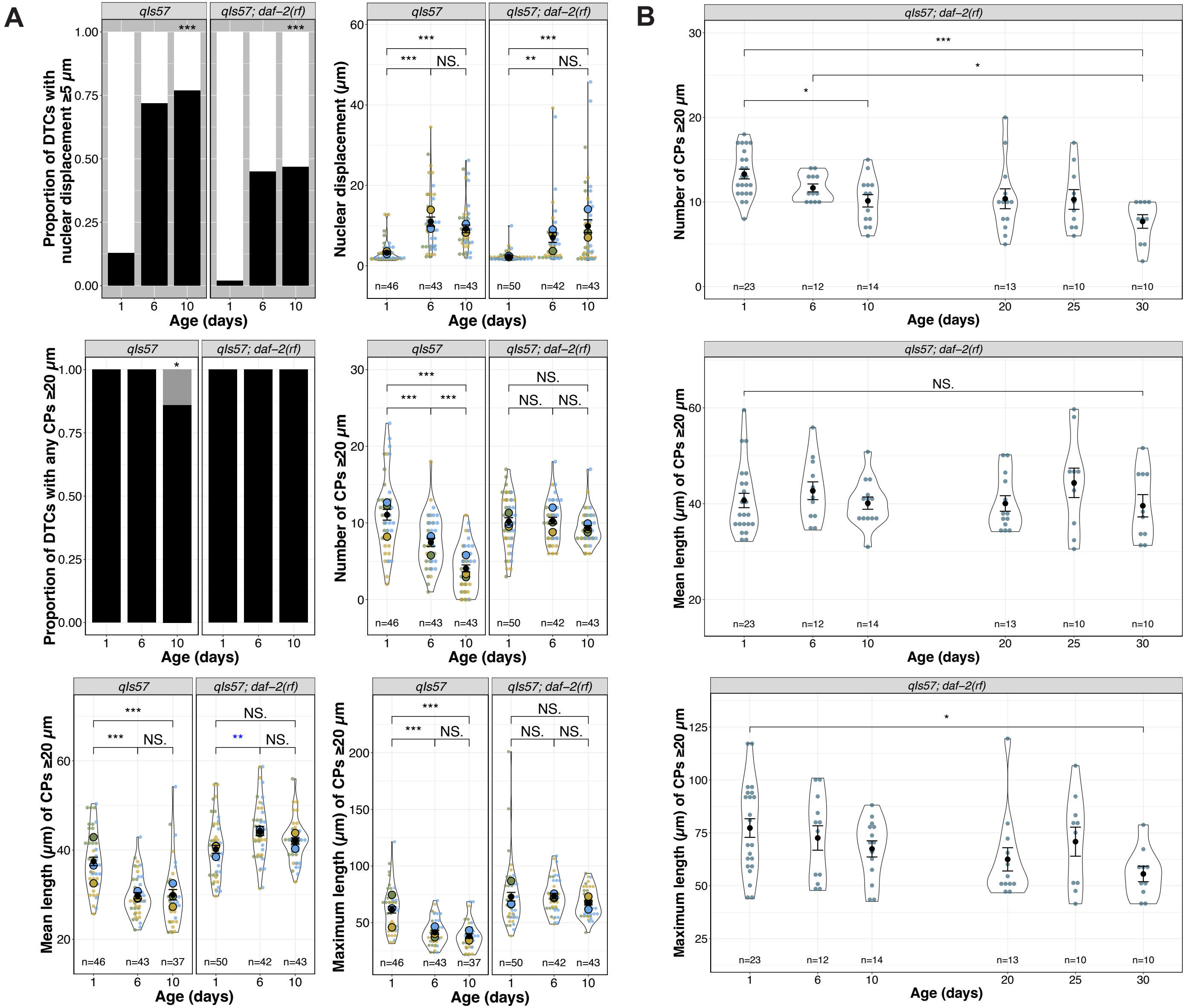

A
B
qIs57
qIs57; daf−2(rf)
Proportion of DTCs with nuclear displacement ≥5 μm
Nuclear displacement (μm)
Number of CPs ≥20 μm
Proportion of DTCs with any CPs ≥20 μm
Mean length (μm) of CPs ≥20 μm
Maximum length (μm) of CPs ≥20 μm
Age (days)
***
**
*
NS.
n=46
n=43
n=37
n=50
n=42
n=23
n=12
n=14
n=13
n=10

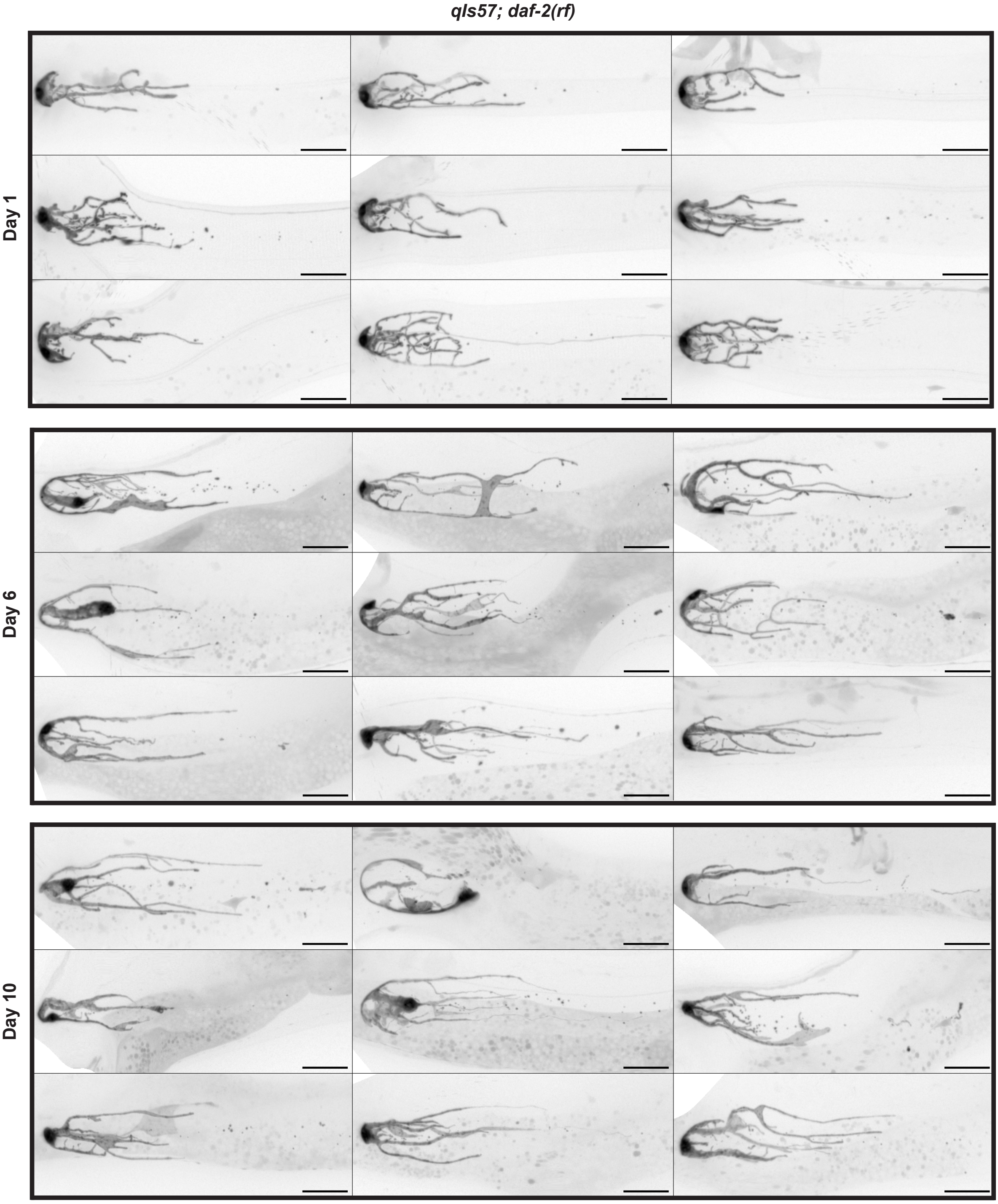

qIs57; daf-2(rf)
Day 1
Day 6
Day 10

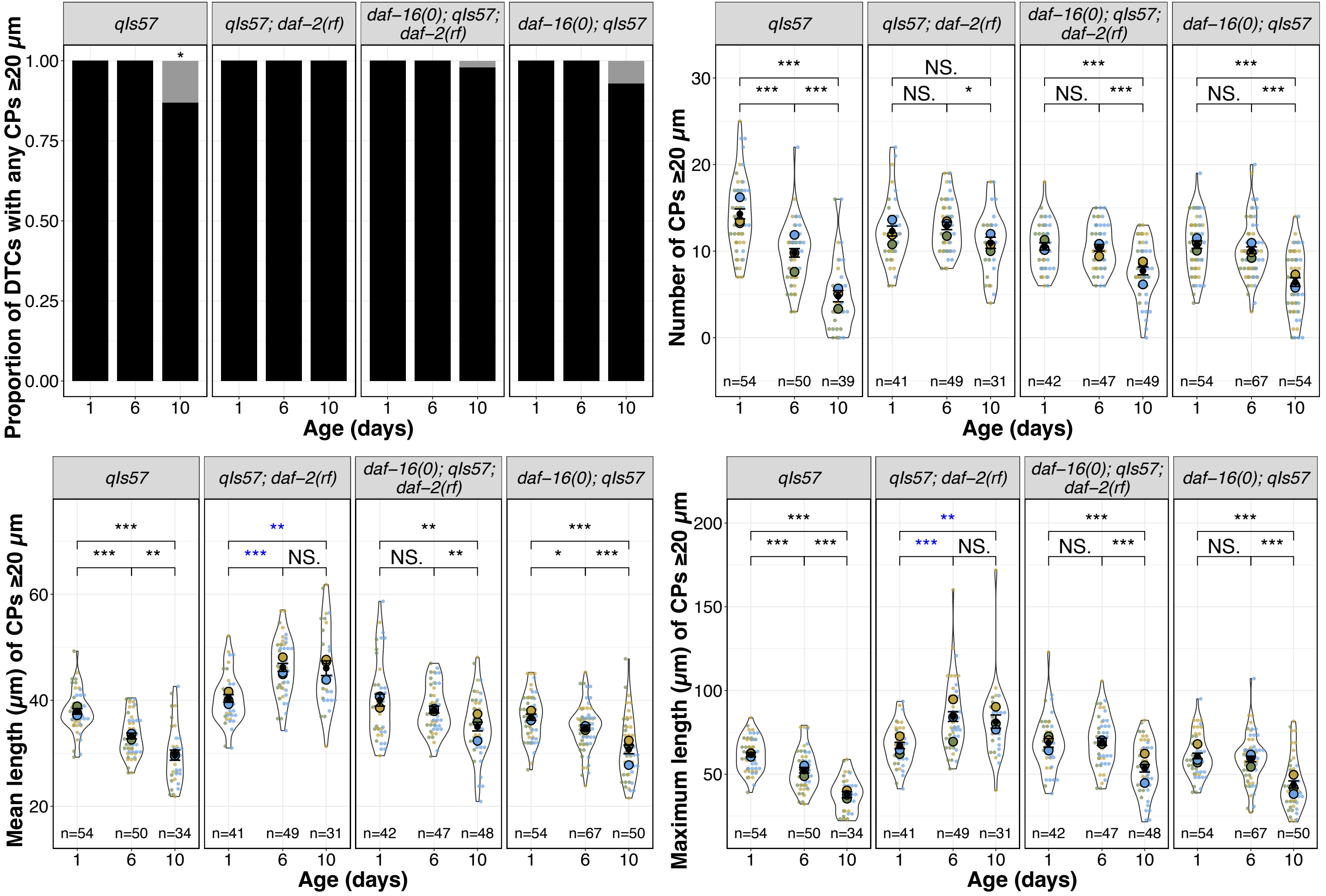

qIs57
qIs57; daf–2(rf)
daf–16(0); qIs57; daf–2(rf)
daf–16(0); qIs57
Proportion of DTCs with any CPs ≥20 μm
Number of CPs ≥20 μm
Mean length (μm) of CPs ≥20 μm
Maximum length (μm) of CPs ≥20 μm
Age (days)
n=54 n=50 n=39
n=41 n=49 n=31
n=42 n=47 n=49
n=54 n=67 n=54
n=54 n=50 n=34
n=42 n=47 n=48
n=54 n=67 n=50
NS.

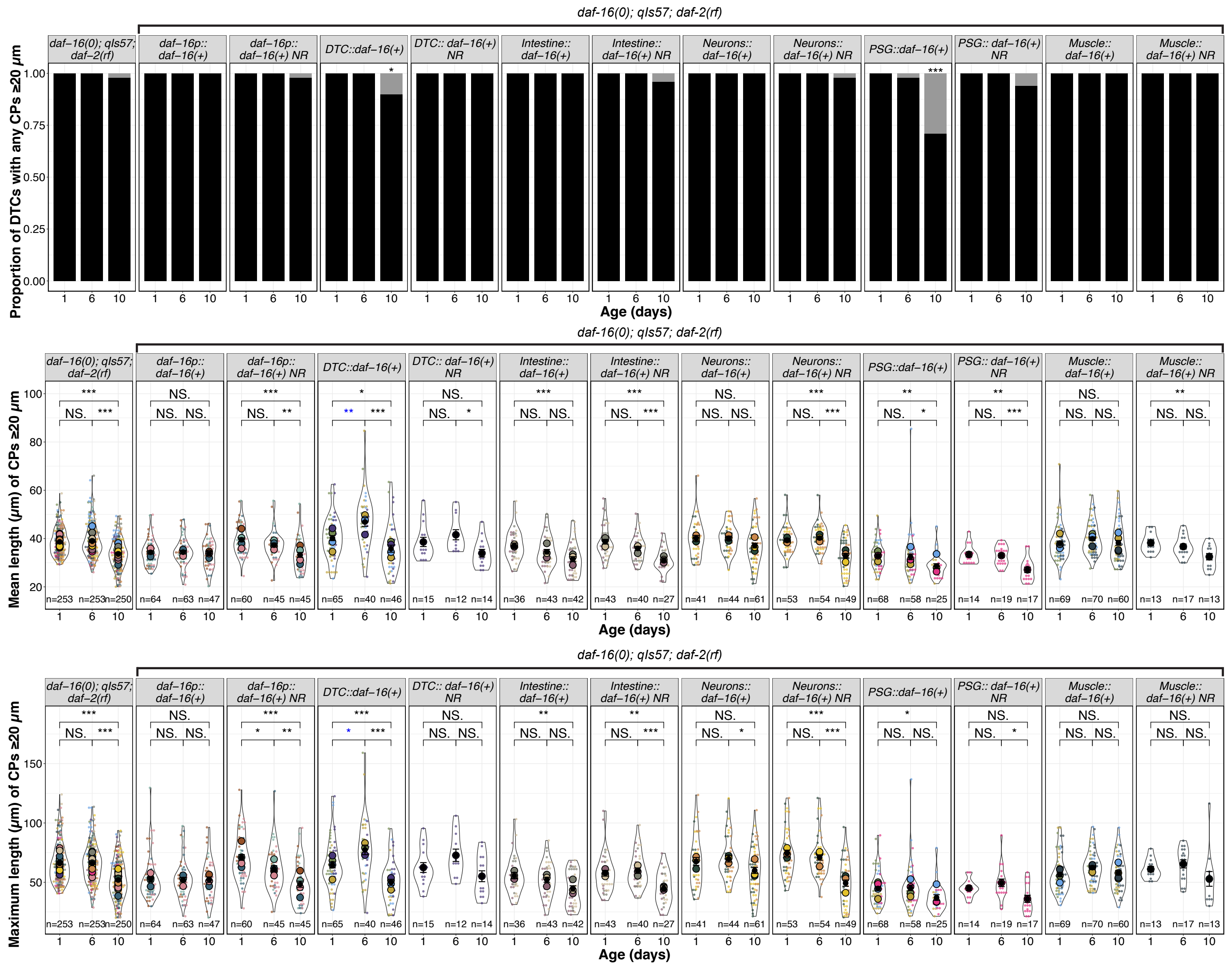
daf-16(0); qIs57; daf-2(rf)
Proportion of DTCs with any CPs ≥20 μm
Mean length (μm) of CPs ≥20 μm
Maximum length (μm) of CPs ≥20 μm
Age (days)
daf–16(0); qIs57; daf–2(rf)
daf–16p:: daf–16(+)
daf–16p:: daf–16(+) NR
DTC::daf–16(+)
DTC:: daf–16(+) NR
Intestine:: daf–16(+)
Intestine:: daf–16(+) NR
Neurons:: daf–16(+)
Neurons:: daf–16(+) NR
PSG::daf–16(+)
PSG:: daf–16(+) NR
Muscle:: daf–16(+)
Muscle:: daf–16(+) NR
n=253 n=253 n=250
n=64 n=63 n=47
n=60 n=45 n=45
n=65 n=40 n=46
n=15 n=12 n=14
n=36 n=43 n=42
n=43 n=40 n=27
n=41 n=44 n=61
n=53 n=54 n=49
n=68 n=58 n=25
n=14 n=19 n=17
n=69 n=70 n=60
n=13 n=17 n=13

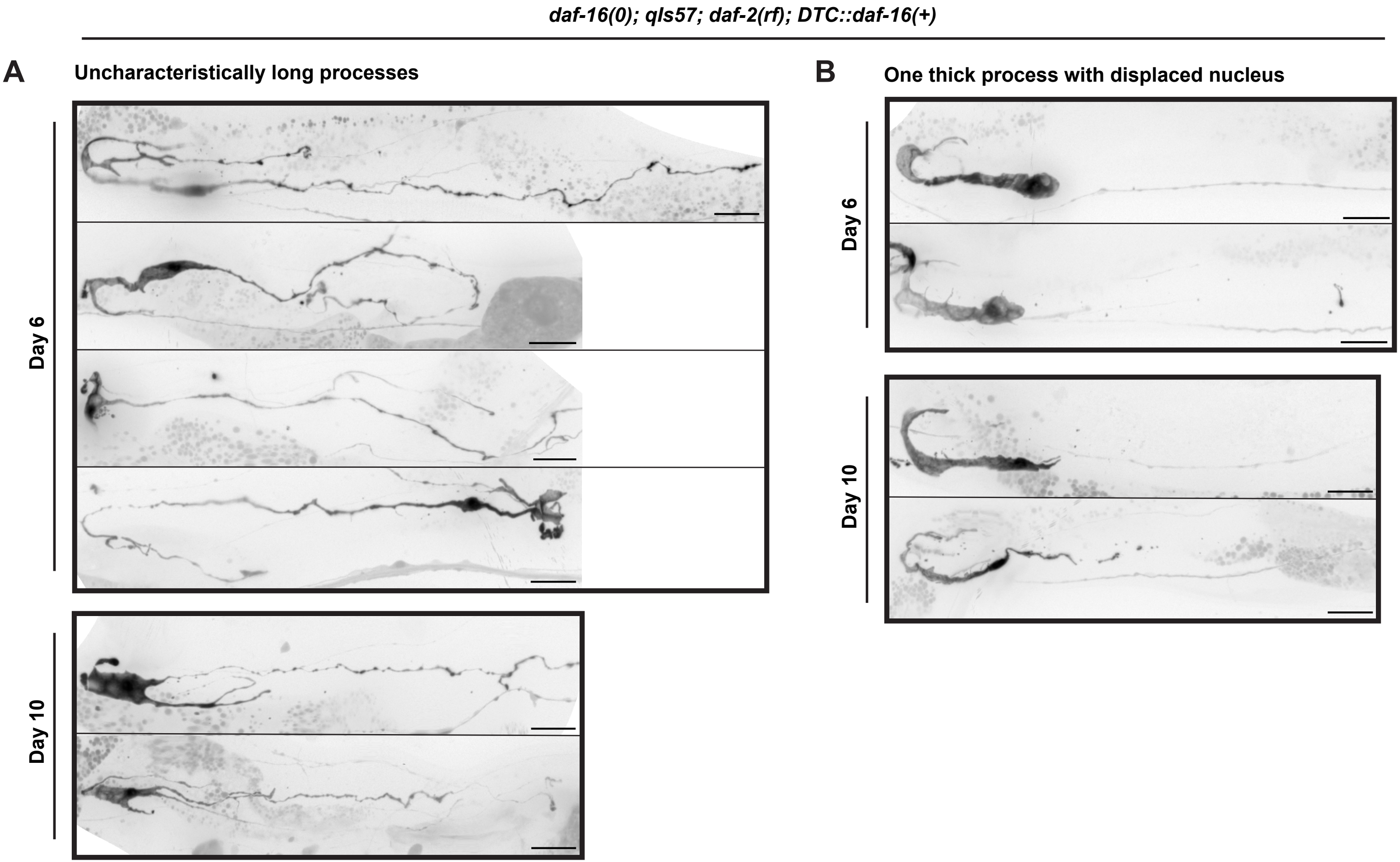

daf-16(0); qIs57; daf-2(rf); DTC::daf-16(+)
A
Uncharacteristically long processes
Day 6
Day 10
B
One thick process with displaced nucleus
Day 6
Day 10

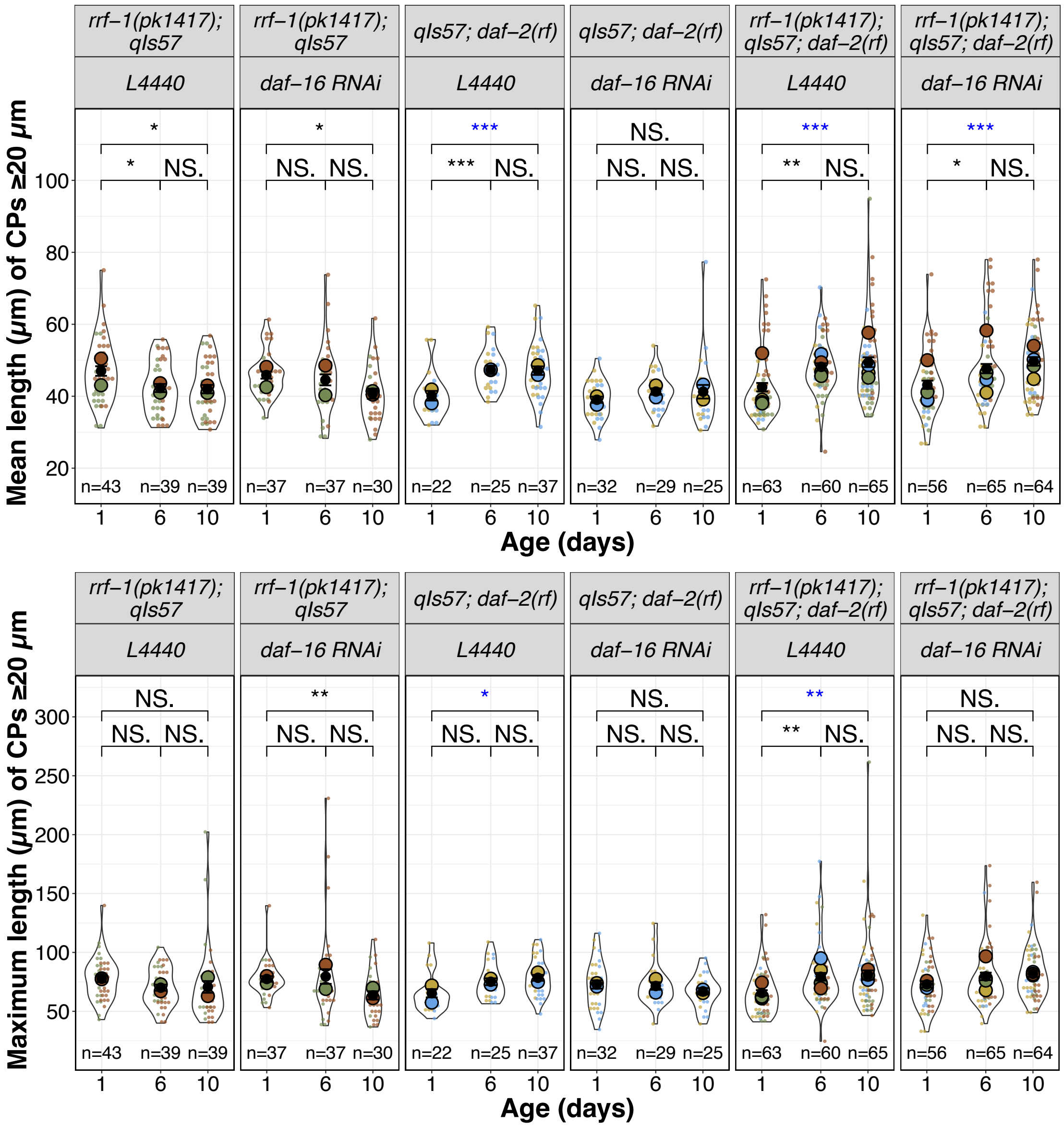

rrf−1(pk1417); qIs57
L4440
rrf−1(pk1417); qIs57
daf−16 RNAi
qIs57; daf−2(rf)
L4440
qIs57; daf−2(rf)
daf−16 RNAi
rrf−1(pk1417); qIs57; daf−2(rf)
L4440
rrf−1(pk1417); qIs57; daf−2(rf)
daf−16 RNAi
Mean length (μm) of CPs ≥20 μm
Age (days)
n=43 n=39 n=39
n=37 n=37 n=30
n=22 n=25 n=37
n=32 n=29 n=25
n=63 n=60 n=65
n=56 n=65 n=64
NS.
Maximum length (μm) of CPs ≥20 μm
Age (days)

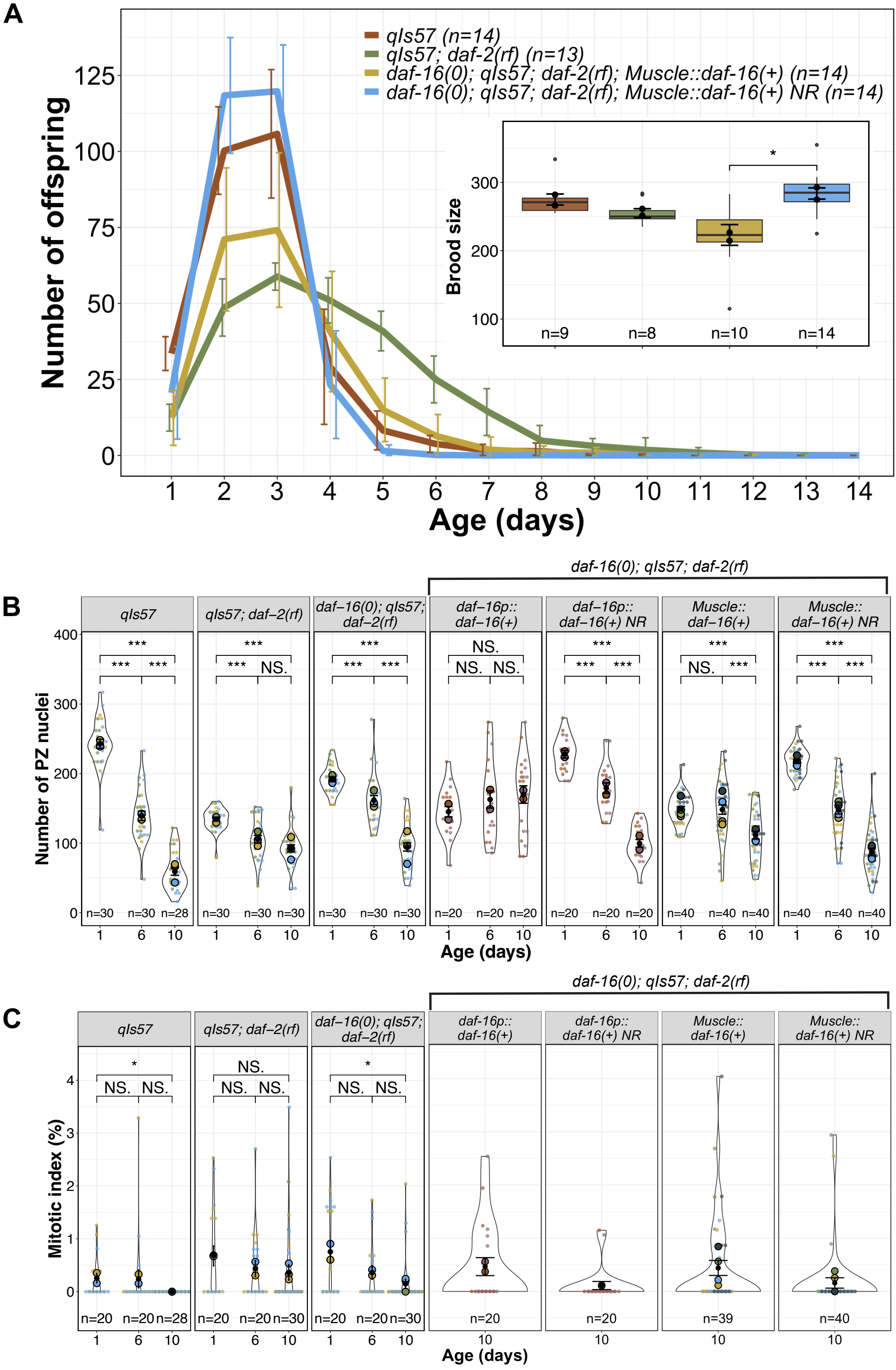
A
qIs57 (n=14)
qIs57; daf-2(rf) (n=13)
daf-16(0); qIs57; daf-2(rf); Muscle::daf-16(+) (n=14)
daf-16(0); qIs57; daf-2(rf); Muscle::daf-16(+) NR (n=14)
Number of offspring
Age (days)
Brood size
n=9
n=8
n=10
n=14
*
B
daf-16(0); qIs57; daf-2(rf)
qIs57
qIs57; daf−2(rf)
daf−16(0); qIs57; daf−2(rf)
daf−16p:: daf−16(+)
daf−16p:: daf−16(+) NR
Muscle:: daf−16(+)
Muscle:: daf−16(+) NR
Number of PZ nuclei
Age (days)
***
NS.
n=30
n=28
n=20
n=40
C
daf-16(0); qIs57; daf-2(rf)
qIs57
qIs57; daf−2(rf)
daf−16(0); qIs57; daf−2(rf)
daf-16p:: daf-16(+)
daf-16p:: daf-16(+) NR
Muscle:: daf-16(+)
Muscle:: daf-16(+) NR
Mitotic index (%)
Age (days)
*
NS.
n=20
n=28
n=30
n=39
n=40

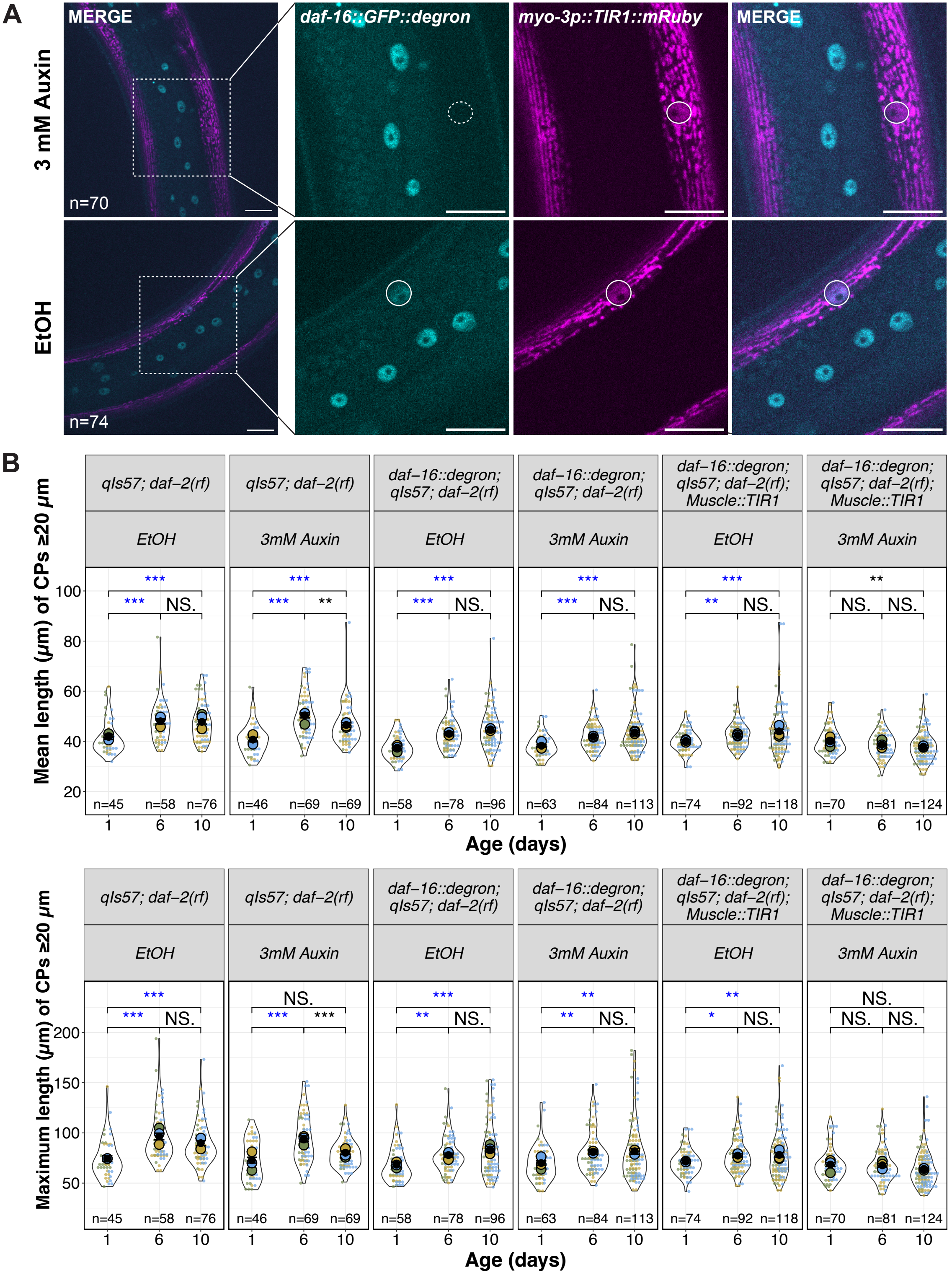

A
3 mM Auxin
EtOH
MERGE
daf-16::GFP::degron
myo-3p::TIR1::mRuby
MERGE
n=70
n=74
B
Mean length (μm) of CPs ≥20 μm
qIs57; daf−2(rf)
daf−16::degron; qIs57; daf−2(rf)
daf−16::degron; qIs57; daf−2(rf); Muscle::TIR1
EtOH
3mM Auxin
n=45 n=58 n=76
n=46 n=69 n=69
n=58 n=78 n=96
n=63 n=84 n=113
n=74 n=92 n=118
n=70 n=81 n=124
Age (days)
Maximum length (μm) of CPs ≥20 μm
NS.

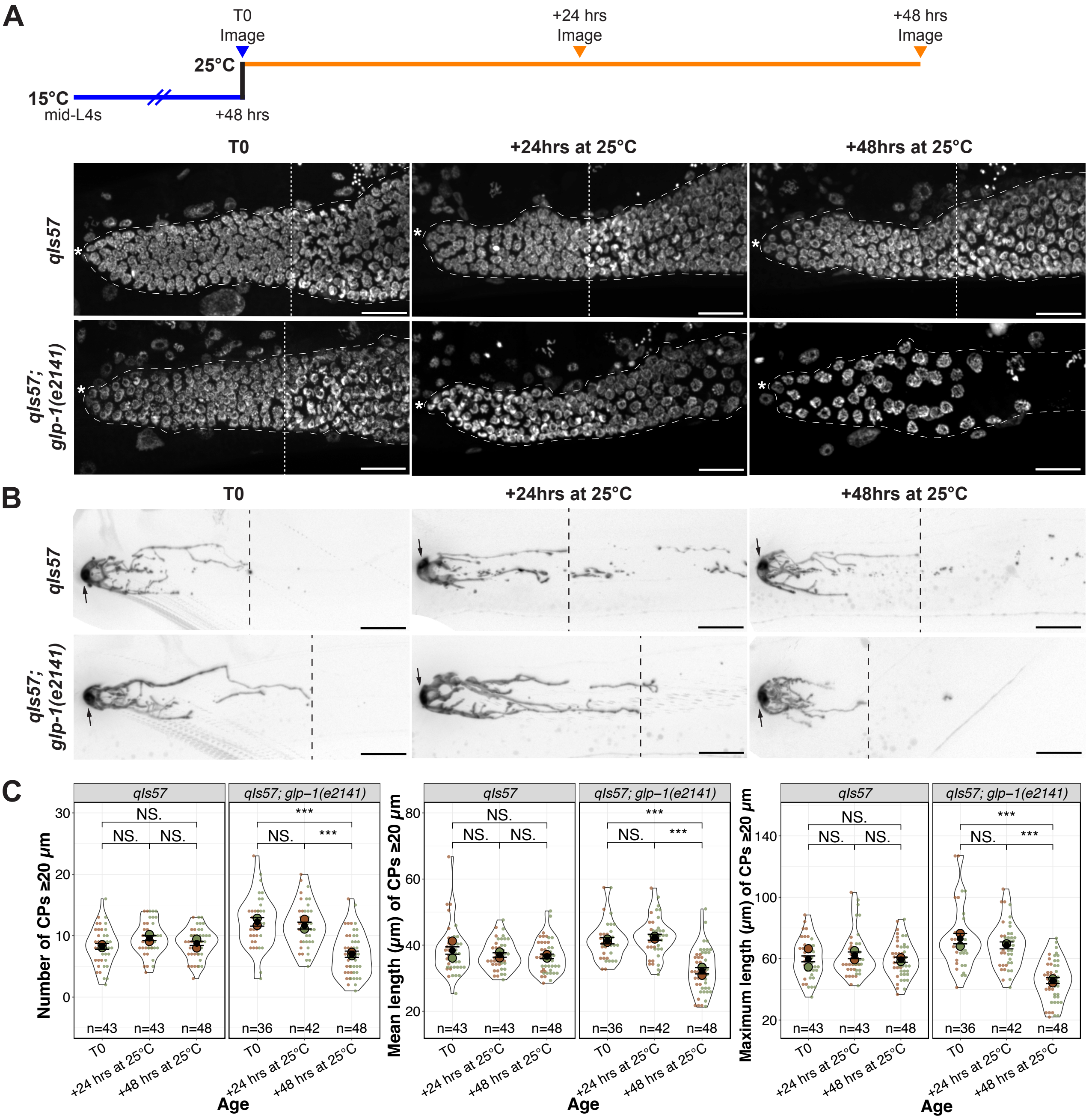

A
T0 Image
+24 hrs Image
+48 hrs Image
25°C
15°C
mid-L4s
+48 hrs
T0
+24hrs at 25°C
+48hrs at 25°C
qIs57
qIs57; glp-1(e2141)
B
C
NS.
***
Number of CPs ≥20 μm
Mean length (μm) of CPs ≥20 μm
Maximum length (μm) of CPs ≥20 μm
n=43
n=48
n=36
n=42
+24 hrs at 25°C
+48 hrs at 25°C
Age

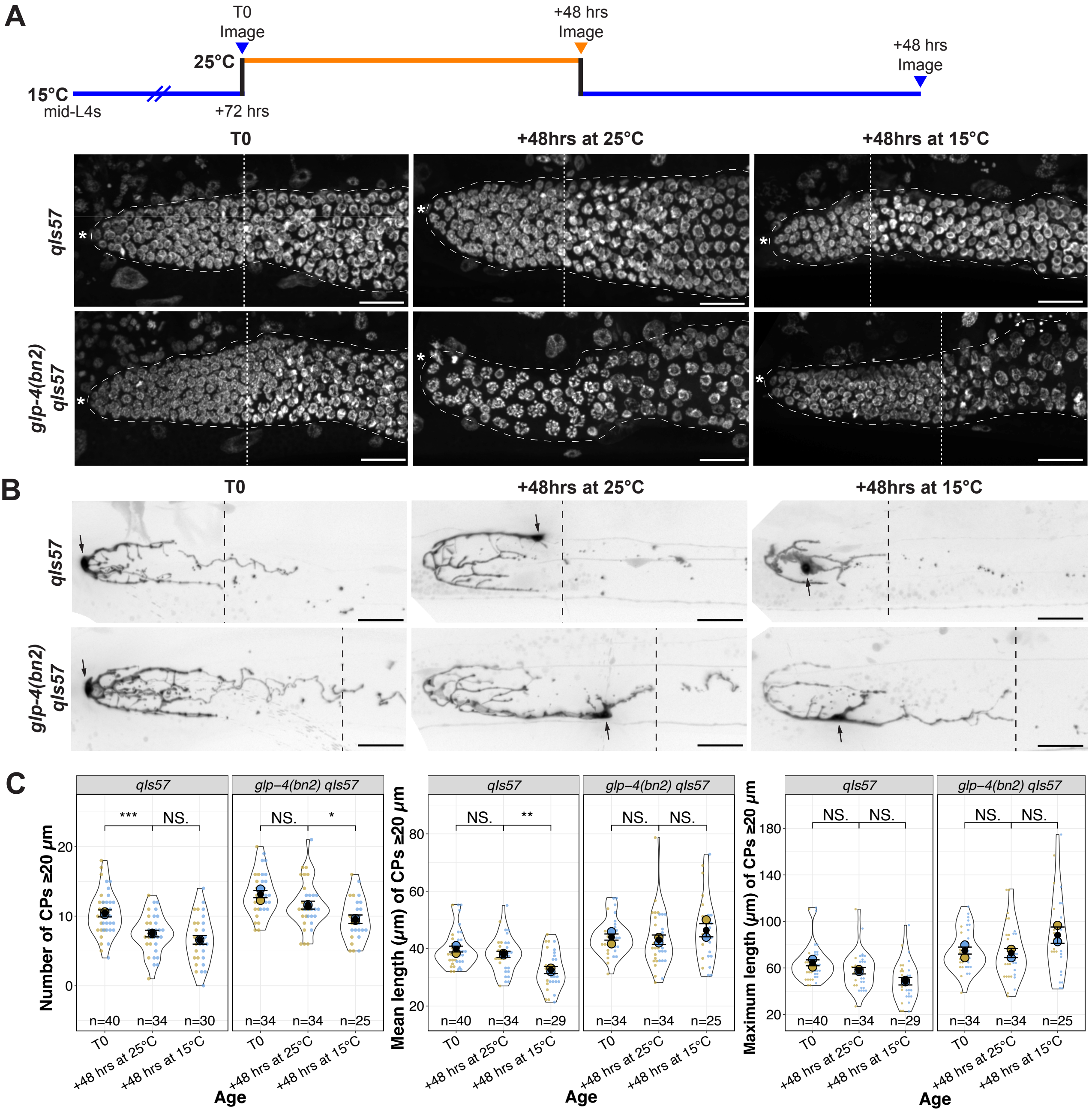

A
T0 Image
+48 hrs Image
+48 hrs Image
25°C
15°C
mid-L4s
+72 hrs
T0
+48hrs at 25°C
+48hrs at 15°C
qIs57
glp-4(bn2) qIs57
B
T0
+48hrs at 25°C
+48hrs at 15°C
qIs57
glp-4(bn2) qIs57
C
qIs57
glp−4(bn2) qIs57
Number of CPs ≥20 μm
Mean length (μm) of CPs ≥20 μm
Maximum length (μm) of CPs ≥20 μm
***
NS.
*
**
n=40
n=34
n=30
n=25
n=29
T0
+48 hrs at 25°C
+48 hrs at 15°C
Age

#### Supplementary Figure Legends

**Figure S1. Comparison of various *lag-2* promoter-driven transgenes**
Representative live worm micrographs of DTCs in strains bearing different *lag-2p*-driven fluorescent proteins at Day 1 and Day 10. Relevant genotypes and features for this study are noted. Scale bars are 20 μm.

**Figure S2. Examples of DTCs in live worms carrying *qIs57* at Day 1, 6, 10 post mid-L4.** Scale bars are 20 μm.

**Figure S3. Age-related DTC parameters and effects of *daf-2(rf)* at Days 1, 6, and 10.**
**(A)** DTC features (from top left to bottom right): proportion of DTCs with nuclear displacement ≥5 μm (black bar), position of nuclear displacement (μm) from the distal end, proportion of DTCs with any CPs ≥20 μm (black bar), number of CPs ≥20 μm per DTC, mean and maximum length of CPs ≥20 μm per DTC. N values for proportion plots are the same as for the "Nuclear Displacement" superplot (top right). For superplots, each dot represents a single DTC (n value shown), colors indicate parallel cohorts, large circles in color are averages for each cohort, and black dot is pooled average for all cohorts shown. N values for mean and maximum length plots reflect only those DTCs with CPs ≥ 20 μm. Day 1 and 10 data are as in Figure 1.
**(B)** Number, mean and maximum length of CPs per DTC in *daf-2(rf)* at indicated ages in days post mid-L4. Day 1, 10, 20 and 30 data are as in Figure 1. **For all panels**, *qIs57* carries *lag-2p::GFP*, and *daf-2(rf)* is *daf-2(e1370).* See methods, Tables S2, S3 for statistics details. For proportion plots, only $p < 0.05$ are indicated; otherwise, NS is "not significant", * $p \geq 0.05$; **$p < 0.01$, *** $p < 0.001$, and blue asterisks indicate a significant increase.

**Figure S4. Examples of DTCs in live worms carrying *qIs57* and *daf-2(e1370)* at Day 1, 6, 10 post mid-L4.** Scale bars are 20 μm.

**Figure S5. Effects of *daf-2* and *daf-16* on DTC process number and length at Days 1, 6, and 10.**
DTC features from top to bottom: proportion of DTCs with any CPs beyond 20 μm, number of CPs ≥20 μm per DTC, mean and maximum length of CPs ≥20 μm per DTC at Day 1, 6, and 10 for the indicated genotypes. N values for proportion plots are as in the Number of CPs plot to the right. For superplots, each dot represents a single DTC n value, colors indicate parallel

cohorts, large circles in color are averages for each cohort, and black dot is pooled average for all cohorts shown. Day 1 and 10 data are as in Figure 2. **For all panels**, *qIs57* carries *lag-2p::GFP*, *daf-16(0)* is *daf-16(mu86),* and *daf-2(rf)* is *daf-2(e1370).* See methods, Tables S2, S3 for statistics details; NS is "not significant", * $p < 0.05$, ** $p < 0.01$, *** $p < 0.001$, and blue asterisks indicate a significant increase.

**Figure S6. Effects of tissue restricted *daf-16a(+)* expression on DTC process length with age.**
Proportion of DTCs with any CPs ≥20 µm, and mean and maximum length of CPs ≥20 µm for indicated genotypes at Day 1, 6 and 10 post mid-L4 plus additional controls. N values for mean and maximum length plots reflect only those DTCs with CPs ≥ 20 µm. NR indicates "non-Rol" siblings from a Rol (array bearing) mother. **In all cases,** *qIs57* carries *lag-2p::GFP*, *daf-16(0)* is *daf-16(mu86),* and *daf-2(rf)* is *daf-2(e1370).* For all superplots, colors indicate cohorts within panels; Day 1 and Day 10 data correspond to main Figure 3. See methods, Tables S2, S3 for statistics details; NS is "not significant", * $p < 0.05$, ** $p < 0.01$, *** $p < 0.001$, and blue asterisks indicate a significant increase.

**Figure S7. Examples of low-penetrance abnormalities in DTC morphology observed in worms over-expressing *daf-16a* in the DTC.** Specific alleles are *daf-16(mu86); qIs57[lag-2p::GFP]; daf-2(e1370); naEx202[lag-2p::daf-16a].* Scale bars are 20 µm.

**Figure S8. Effects of *daf-16 RNAi* in *daf-2(e1370)* on Day 1, 6, and 10 on mean and maximum length of CPs.** See methods, Tables S2, S3 for statistics details; NS is "not significant", * $p < 0.05$, ** $p < 0.01$, *** $p < 0.001$, and blue asterisks indicate a significant increase. Day 1 and Day 10 data shown are as in main Figure 3B.

**Figure S9. The *daf-2(rf)* age-related extended reproductive period, delay in the PZ pool decline and elevated mitotic index are variably affected by muscle-expressed *daf-16a(+).***
**(A)** Muscle-expressed *daf-16a(+)* does not prolong the self-fertile reproductive period seen in *daf-2(rf)*. Offspring timeline and brood size analysis. Number of offspring produced by self-fertile hermaphrodites over time in indicated genotypes. The number of offspring produced on specific days in *daf-16(0);daf-2(rf)* worms bearing muscle-expressed *daf-16a(+)* is significantly different from wild type only for Day 1 and 2, from non-roller (NR) siblings on Day 2 and 3, and from *daf-2(rf)* on Days 5-7. In addition, wild type and NR differ on Day 6 only. All other comparisons are

not significant, see Table S3 for p-values. Progeny counts from individuals that died during the analysis are included. **Inset:** Total brood count box plots indicate the median (bold line) and interquartile ranges. Solid black dots indicate replicate means. Outliers are shown when they fall outside 1.5x of the interquartile range. These counts exclude individuals that died prior to a zero-offspring day. The only significant difference among all pair-wise comparisons is the one indicated.

**(B) Progenitor zone counts at Day 1, 6, and 10.** Day 1 and Day 10 data correspond to main Figure 4. Additional controls: NR indicates “non-Rol” siblings from a Rol (array bearing) mother.

**(C) Mitotic index at at Day 1, 6, and 10 or Day 10** in the indicated genotypes. Day 10 data correspond to main Figure 4.

**For all panels,** *qIs57* carries *lag-2p::GFP*, *daf-16(0)* is *daf-16(mu86),* and *daf-2(rf)* is *daf-2(e1370).* For all superplots, colors indicate cohorts within panels. See methods, Tables S2, S3 for statistics details; NS is “not significant”, * $p < 0.05$, ** $p < 0.01$, *** $p < 0.001$.

**Figure S10. Effects of body-wall muscle depletion of *daf-16* on DTC process length with age. (A)** Representative examples of DAF-16::GFP in the body wall muscle upon exposure to auxin versus the vehicle control (EtOH). The expression of DAF-16::GFP was visually assessed in all worms in which the DTC was measured (n = 70 and 74). **(B)** Mean and maximum length of CPs ≥20 µm for conditions indicated where *daf-16::degron* is *hq389[daf-16::GFP::degron]* and *Muscle::TIR1* is *emcSi71[myo-3p::TIR-1::mRuby]*, including EtOH controls. For all superplots, colors indicate cohorts within panels; Day 1 and Day 10 data correspond to main Figure 3. See methods, Tables S2, S3 for statistics details; NS is “not significant”, * $p < 0.05$, ** $p < 0.01$, *** $p < 0.001$, and blue asterisks indicate a significant increase.

**Figure S11. Premature decline of DTC processes upon differentiation of underlying germ cells after shift of *glp-1(e2141ts)* to the restrictive temperature.** Initial and 48-hour timepoints (images and graphs) correspond to main Figure 5. **(A)** Experimental design and images of DAPI-stained gonads in control and *glp-1(e2141ts)* worms at indicated timepoints. White dotted lines indicate the proximal border of the PZ. Asterisks indicate the distal end. **(B)** Examples of DTC morphology in worms scored in parallel to those shown in panel (A). Black dashed lines indicate the longest CP. Arrows indicate the DTC nucleus. **(A, B)** Scale bars are 20 µm. (C) Quantification of number, mean and maximum lengths of CPs ≥20 µm in control and *glp-1(e2141ts)*. Colors indicate cohorts within panels. See methods, Tables S2, S3 for statistics details; NS is "not significant", * $p < 0.05$, ** $p < 0.01$, *** $p < 0.001$.

**Figure S12. DTC processes do not decline upon *glp-4(bn2ts)* shift to restrictive temperature and remain intact after return to permissive temperature.** Initial and 48-hour timepoints (images and graphs) correspond to main Figure 5. Full analysis includes shift back to 15ºC where germ cells return to a more normal appearance and resume divisions. **(A)** Experimental design and images of DAPI-stained gonads in control and *glp-4(bn2ts)* worms at indicated timepoints. White dotted lines indicate the proximal border of the PZ. Asterisks indicate the distal end. **(B)** Examples of DTC morphology in worms scored in parallel to those shown in panel (A). Black dashed lines indicate the longest CP. Arrows indicate the DTC nucleus. **(A, B)** Scale bars are 20 μm. (C) Quantification of number, mean and maximum lengths of CPs ≥20 μm in control and *glp-4(bn2ts)*. See methods, Tables S2, S3 for statistics details; NS is "not significant", * $p < 0.05$, ** $p < 0.01$, *** $p < 0.001$.

**Table S1. Strains, oligonucleotides, and other reagents used in this study.**

**Table S2. Mean, standard error/deviation, and n values.** Tabs contain values for each main and corresponding supplemental data figures, where applicable.

**Table S3. Genotypes compared, statistical tests, and exact p values.** Tabs contain values for each main and supplemental data figure.

### Table S1

| Strains | | |
|---|---|---|
| ***C. elegans* Strains** | **Genotype** | **References** |
| N2 | wild type (Bristol) | Brenner, 1974 |
| CB1370 | *daf-2(e1370) III* | Riddle *et al.,* 1981 |
| CF1442 | *daf-16(mu86) I; daf-2(e1370) III; muEx169[unc-119p::GFP::daf-16 +* pRF4*(rol-6(su1006))]* | Libina *et al.* , 2003 |
| CF1449 | *daf-16(mu86) I; daf-2(e1370) III; muEx176[daf-16p::GFP::daf-16 +* pRF4*(rol-6(su1006))]* | Libina *et al.,* 2003 |
| CF1514 | *daf-16(mu86) I; daf-2(e1370) III; muEx211[* pNL213*(ges-1p::GFP::daf-16) +* pRF4*(rol-6(su1006))]* | Libina *et al.* , 2003 |
| CF1515 | *daf-16(mu86) I; daf-2(e1370) III; muEx212[* pNL212*(myo-3p::GFP::daf-16) +* pRF4*(rol-6(su1006))]* | Libina *et al.,* 2003 |
| GC832 | *glp-1(e2141) III* | Priess *et al.* , 1987; Hutter and Schnabel, 1994;<br>Dalfó *et al* ., 2010 |
| GC1019 | *rrf-1(pk1417) I; daf-2(e1370) III* | Michaelson *et al.,* 2010 |
| GC1109 | *daf-16(m26) I; daf-2(e1370) III; naEx202[* pGC461*(lag-2p::daf-16::GFP) +* pRF4*(rol-6(su1006))]* | Michaelson *et al.,* 2010 |
| GC1172 | *xnSi1[mex-5p::GFP::PHdomain::nos-2 3'UTR] II ; naIs37[* pGC457*(lag-2p::PHdomain::mCherry-unc-119(+))]* | Chihara & Nance, 2012;<br>Pekar *et al.,* 2017 |
| GC1285 | *daf-16(m26) I; daf-2(e1370) III; naEx239[* pGC629*(fos-1p::GFP::daf-16) +* pRF4*(rol-6(su1006))]* | Qin & Hubbard, 2015 |
| GC1332 | *daf-16(mu86) I; daf-2(e1370) III* | Qin & Hubbard, 2015 |
| GC1412 | *naSi8[* pGC680*(lag-2p(3kb)::GFP-PH::let-858(3'))] II* | Pekar *et al.,* 2017 |
| GC1593 | *qIs57[pJK590(lag-2p::GFP)] II; glp-1(e2141) III*<br><br>N2 males were crossed with JK2869 hermaphrodites, and the resulting heterozygous males were crossed with GC832 hermaphrodites. *qIs57* was followed visually. g*lp-1(e2141)* was confirmed by DNA sequence analysis. | This study |
| GC1607 | *qIs57[* pJK590*(lag-2p::GFP)] II; daf-2(e1370) III*<br><br>JK2869 males were crossed with CB1370 hermaphrodites, and *qIs57* was followed visually. *daf-2(e1370)* was followed by the Daf-c phenotype and subsequent genotyping by DNA sequence analysis. | This study |
| GC1616 | *daf-16(mu86) I; qIs57[* pJK590*(lag-2p::GFP)] II; daf-2(e1370) III*<br><br>GC1332 males were crossed with GC1607 hermaphrodites, and *qIs57* was followed visually. The presence of both *daf-16(mu86)* and *daf-2(e1370)* were followed by monitoring the Daf-c phenotype and subsequent DNA sequence analysis. | This study |
| GC1676 | *daf-16(mu86) I; qIs57[* pJK590*(lag-2p::GFP)] II; daf-2(e1370) III; naEx202[* pGC461*(lag-2p::daf-16::GFP) +* pRF4*(rol-6(su1006))]*<br><br>GC1616 males were crossed with GC1109 array-bearing (Rol) hermaphrodites, and *qIs57* was followed visually. *daf-16(mu86)* was followed by DNA sequence analysis. DTC-specific expression of *daf-16a(+)* was confirmed visually by presence of GFP in DTC nuclei and some neuronal nuclei. | This study |
| GC1682 | *daf-16(mu86) I; qIs57[* pJK590*(lag-2p::GFP)] II; daf-2(e1370) III; muEx212[* pNL212*(myo-3p::GFP::daf-16) +* pRF4*(rol-6(su1006))]*<br><br>GC1616 males were crossed with CF1515 array-bearing (Rol) hermaphrodites, and *qIs57* was followed visually. *daf-16(mu86)* was followed by DNA sequence analysis. Muscle-specific expression of daf-16a(+) was confirmed by presence of GFP in muscle nuclei. | This study |

| | | |
|---|---|---|
| GC1683 | *daf-16(mu86) I; qIs57[*pJK590*(lag-2p::GFP)] II; daf-2(e1370) III; naEx239[*pGC629*(fos-1p::GFP::daf-16)* + pRF4*(rol-6(su1006))]*<br><br>GC1616 males were crossed with GC1285 array-bearing (Rol) hermaphrodites, and *qIs57* was followed visually. *daf-16(mu86)* was followed by DNA sequence analysis. PSG-specific expression of *daf-16a(+)* confirmed visuallly by presence of GFP in PSG nuclei. | This study |
| GC1691 | *daf-16(mu86) I; qIs57[pJK590(lag-2p::GFP)] II; daf-2(e1370) III; muEx176[daf-16p::GFP::daf-16a* + pRF4*(rol-6(su1006))]*<br><br>GC1616 males were crossed with CF1449 array-bearing (Rol) hermaphrodites, and *qIs57* was followed visually. *daf-16(mu86)* was followed by DNA sequence analysis. Expression of *daf-16a(+)* was confirmed visually by the presence of GFP in multiple tissues. | This study |
| GC1692 | *daf-16(mu86) I; qIs57[*pJK590*(lag-2p::GFP)] II; daf-2(e1370) III; muEx211[*pNL213*(ges-1p::GFP::daf-16)* + pRF4*(rol-6(su1006))]*<br><br>GC1616 males were crossed with CF1514 array-bearing (Rol) hermaphrodites, and *qIs57* was followed visually. *daf-16(mu86)* was followed by DNA sequence analysis. Intestine-specific expression of *daf-16a(+)* was confirmed visually by presence of GFP in intestinal nuclei. | This study |
| GC1696 | *daf-16(mu86) I; qIs57[*pJK590*(lag-2p::GFP)] II; daf-2(e1370) III; muEx169[unc-119p::GFP::daf-16* + pRF4*(rol-6(su1006))]*<br><br>GC1616 males were crossed to CF1442 array-bearing (Rol) hermaphrodites, and *qIs57* was followed visually. *daf-16(mu86)* was followed by DNA sequence analysis. Neuron-specific expression of *daf-16a(+)* was confirmed visually by presence of GFP in neuronal nuclei. | This study |
| GC1701 | *glp-4(bn2) qIs57[pJK590(lag-2p::GFP)] II* | Tolkin *et al.,* 2024 |
| GC1748 | *rrf-1(pk1417) I; qIs57[*pJK590(*lag-2p::GFP)] II*<br><br>JK4143 males were crossed with PD8488 hermaphrodites. *qIs57* was followed visually and *rrf-1(pk1417)* was confirmed by DNA sequence analysis. | This study |
| GC1778 | *daf-16(mu86) I; qIs57*[pJK590*(lag-2p::GFP)*] *II*<br><br>JK2869 males were crossed with GC1616 hermaphrodites. *daf-2(e1370)* was selected against by absence of the Daf-c phenotype. Both *daf-2(e1370)* and *daf-16(mu86)* were subsequently confirmed by DNA sequence analysis. | This study |
| GC1817 | *daf-16(hq389[daf-16::GFP::degron]) I; qIs57[*pJK590*(lag-2p::GFP)] II; daf-2(e1370) III; emcSi71[myo-3p::TIR-1::mRuby] (IV: -0.05)*<br><br>HAL230 males were crossed with GC1819 hermaphrodites, and *qIs57* was followed visually. *daf-2(e1370)* was followed by the Daf-c phenotype and subsequent genotyping by DNA sequence analysis. *daf-16::GFP::degron* was confirmed visually by the presence of GFP in the nuclei of multiple tissues and by subsequent genotyping by DNA sequence analysis. *myo-3p::TIR1::mRuby* was confirmed visually by the presence of red fluoresence in muscle nuclei and by subsequent genotyping by DNA sequence analysis. | This study |
| GC1819 | *daf-16(hq389[daf-16::GFP::degron]) I; qIs57[*pJK590*(lag-2p::GFP)] II; daf-2(e1370) III*<br><br>GC1607 males were crossed with MQD2499 hermaphrodites, and *qIs57* was followed visually. *daf-2(e1370)* was followed by the Daf-c phenotype. *daf-16::GFP::degron* was confirmed visually by the presence of GFP in the nuclei of multiple tissues. *myo-3p::TIR1::mRuby* was selected against by the absence of red fluorescence in the muscle. All three alleles were confirmed by subsequent genotyping by DNA sequence analysis. | This study |
| GC1912 | *rrf-1(pk1417) I; qIs57[*pJK590*(lag-2p::GFP)] II; daf-2(e1370) III*<br><br>GC1607 males were crossed with GC1019 hermaphrodites, and *qIs57* was followed visually. *daf-2(e1370)* was followed by the Daf-c phenotype and by subsequent genotyping by DNA sequence analysis. *rrf-1(pk1417)* was also confirmed by DNA sequence analysis. | This study |
| HAL230 | *unc-119(ed3) III; emcSi71[myo-3p::TIR1::mRuby] (IV: -0.05)* | Sabatella *et al.,* 2021 |

| | | |
|---|---|---|
| JK2869 | *qIs57[* pJK590*(lag-2p::GFP)] II* | Siegfried ***et al.,*** 2004 |
| JK4143 | *qIs57[pJK590(lag-2p::GFP)] II; rde-1(ne219) V; qIs140[lag-2p::RDE-1 +* pRF4(rol-6(su1006))] | Martynovsky ***et al.,*** 2012 |
| JK4472 | *qIs154[lag-2p::MYR::tdTomato + ttx-3p::GFP] V* | Byrd ***et al.,*** 2014 |
| MQD2499 | *daf-16(hq389[daf-16::GFP::degron]) I; hqSi10[myo-3p::TIR1::mRuby::unc-54 3' UTR + Cbr-unc-119(+)] II; daf-2(e1370) unc-119(ed3) III* | Zhang ***et al.,*** 2022 |
| PD8488 | *rrf-1(pk1417) I* | Miller ***et al.,*** 2003 |
| **Bacterial Strains** | **Genotype** | **References** |
| *E. coli* HT115(DE3) | [F-, mcrA, mcrB, IN(rrnD-rrnE)1, rnc14::Tn10(DE3 lysogen: lacUV5 promoter -T7 polymerase] | Timmons ***et al.,*** 2001 |
| *E. coli* HT115(DE3) L4440 | HT115(DE3) bacteria bearing the L4440 "empty vector" plasmid | Timmons ***et al.,*** 2001 |
| *E. coli* HT115(DE3) 10020-B-8 | HT115(DE3) bearing a plasmid that produces dsRNA corresponding to the *daf-16* cDNA | Rual ***et al.,*** 2004 |
| *E. coli* OP50 | *E. coli* Uracil auxotroph | CGC |

#### Oligonucleotides

| **Name** | **Sequence** | **Gene** |
|---|---|---|
| GCo969 | GCCGCACAGATTTGTGATGGTATGGCG | F primer for *daf-2(e1370)* |
| GCo970 | TCATCAAGATCCAGTGCTTCTGAATCG | R primer for *daf-2(e1370)* |
| GCo1217 | CTCGCAAATGGATATCCCGG | F primer for *glp-1(e2141)* |
| GCo1354 | CATCCTTCCTTAGCGGCAAGC | R primer for *glp-1(e2141)* |
| GCo1624 | CGAATCAGTTTTTTTTCTTCACTCGCC | F primer for *daf-16(mu86)* |
| GCo1625 | CGTGAGAAATCGTTTGAATCGATCCGGC | R primer for *daf-16(mu86)* |
| GCo1971 | TAGCTATCAAATCCAATACTGGGAG | F primer for *rrf-1(pk1417)* |
| GCo1976 | GTTGTGGGAATACTACTACATCAC | R primer for *rrf-1(pk1417)* |
| GCo2949 | CAGCCTATACGGGAGCAATGAGCA | F primer for *daf-16(m26)* |
| GCo2950 | GCGTTCCTATACGAATGAAACCAACTGG | R primer for *daf-16(m26)* |
| GCo3177 | CCTCTACTTCTGCCGTCAAATGACCAACG | F primer for *TIR1* |
| GCo3178 | ACAGATCCCTCAAGAACAACCTTCATGCG | R primer for *TIR1* |
| GCo3183 | TTCAGAATGCCACTACCGGGTGCCTAT | F primer for *daf-16::degron* |
| GCo3184 | TGTTTGCAGTTCTAAGGCCCTTATTGGA | R primer for *daf-16::degron* |
| GCo3225 | TGGCTCATTCCTTTATCGTCATACA | F primer for *rde-1(ne300)* |
| GCo3226 | CGAGTATCAATCCAGGTGGAACTAT | R primer for *rde-1(ne300)* |

#### Reagents

| **Name** | **Source** | **Identifier** |
|---|---|---|
| Indole-3-acetic acid (Auxin) | Thermo Scientific | A10556.14 |
| VECTASHIELD® Antifade Mounting Medium, With DAPI, Liquid | Vector Laboratories | H-1200-10 |
| β-Lactose | Sigma-Aldrich | L3750 |

**Supplementary references**

**Brenner, S.** (1974). The genetics of Caenorhabditis elegans. *Genetics* **77**, 71-94.

**Byrd, D. T., Knobel, K., Affeldt, K., Crittenden, S. L. and Kimble, J.** (2014). A DTC niche plexus surrounds the germline stem cell pool in Caenorhabditis elegans. *PLoS One* **9**, e88372.

**Chihara, D. and Nance, J.** (2012). An E-cadherin-mediated hitchhiking mechanism for C. elegans germ cell internalization during gastrulation. *Development*.

**Dalfo, D., Priess, J. R., Schnabel, R. and Hubbard, E. J. A.** (2010). glp-1(e2141) sequence correction. In *Worm Breeder's Gazette*.

**Hutter, H. and Schnabel, R.** (1994). glp-1 and inductions establishing embryonic axes in C. elegans. *Development* **120**, 2051-2064.

**Libina, N., Berman, J. R. and Kenyon, C.** (2003). Tissue-specific activities of C. elegans DAF-16 in the regulation of lifespan. *Cell* **115**, 489-502.

**Martynovsky, M., Wong, M. C., Byrd, D. T., Kimble, J., & Schwarzbauer, J. E.** (2012). mig-38, a novel gene that regulates distal tip cell turning during gonadogenesis in C. elegans hermaphrodites. *Developmental Biology*, **368**, 404–414.

**Michaelson, D., Korta, D. Z., Capua, Y. and Hubbard, E. J. A.** (2010). Insulin signaling promotes germline proliferation in C. elegans. *Development (Cambridge, England)* **137**, 671-680.

**Miller, M. A., Ruest, P. J., Kosinski, M., Hanks, S. K. and Greenstein, D.** (2003). An Eph receptor sperm-sensing control mechanism for oocyte meiotic maturation in Caenorhabditis elegans. *Genes & development* **17**, 187-200.

**Pekar, O., Ow, M. C., Hui, K. Y., Noyes, M. B., Hall, S. E. and Hubbard, E. J. A.** (2017). Linking the environment, DAF-7/TGFbeta signaling and LAG-2/DSL ligand expression in the germline stem cell niche. *Development* **144**, 2896-2906.

**Priess, J. R., Schnabel, H. and Schnabel, R.** (1987). The glp-1 locus and cellular interactions in early C. elegans embryos. *Cell* **51**, 601-611.

**Qin, Z. and Hubbard, E. J. A.** (2015). Non-autonomous DAF-16/FOXO activity antagonizes age-related loss of C. elegans germline stem/progenitor cells. *Nature Communications* **6**, 7107.

**Riddle, D. L., Swanson, M. M. and Albert, P. S.** (1981). Interacting genes in nematode dauer larva formation. *Nature* **290**, 668-671.

**Rual, J. F., Ceron, J., Koreth, J., Hao, T., Nicot, A. S., Hirozane-Kishikawa, T., Vandenhaute, J., Orkin, S. H., Hill, D. E., van den Heuvel, S., et al.** (2004). Toward improving Caenorhabditis elegans phenome mapping with an ORFeome-based RNAi library. *Genome Res* **14**, 2162-2168.

**Sabatella, M., Thijssen, K. L., Davo-Martinez, C., Vermeulen, W. and Lans, H.** (2021). Tissue-Specific DNA Repair Activity of ERCC-1/XPF-1. *Cell Rep* **34**, 108608.

**Siegfried, K. R., Kidd, A. R., 3rd, Chesney, M. A. and Kimble, J.** (2004). The sys-1 and sys-3 genes cooperate with Wnt signaling to establish the proximal-distal axis of the Caenorhabditis elegans gonad. *Genetics* **166**, 171-186.

**Timmons, L., Court, D. L. and Fire, A.** (2001). Ingestion of bacterially expressed dsRNAs can produce specific and potent genetic interference in Caenorhabditis elegans. *Gene* **263**, 103-112.

**Tolkin, T., Burnett, J., & Hubbard, E. J. A.** (2024). A role for organ level dynamics in morphogenesis of the C. elegans hermaphrodite distal tip cell. *Development (Cambridge, England)*, **151**.

**Zhang, Y. P., Zhang, W. H., Zhang, P., Li, Q., Sun, Y., Wang, J. W., Zhang, S. O., Cai, T., Zhan, C. and Dong, M. Q.** (2022). Intestine-specific removal of DAF-2 nearly doubles lifespan in Caenorhabditis elegans with little fitness cost. *Nat Commun* **13**, 6339.
